# The Role of Training Paradigms in Shaping Connectivity and Dynamics: A Comparison of Gradient-Based and Evolutionary Methods in Recurrent Neural Network Models

**DOI:** 10.64898/2026.07.09.737022

**Authors:** Ivyer Mingwei Qu, Jiayue Dora Li, Yuqing Zhu

## Abstract

Recurrent neural networks (RNNs) trained with backpropagation through time (BPTT) use gradients and global error signals to solve tasks, while evolutionary algorithms (EAs) offer an alternative solution through gradient-free optimization. Both classes of methods can solve the same tasks by modifying network weights, but it remains unclear how the choice of training paradigm biases the final connectivity structure and resulting dynamics. When drawing conclusions from task-trained RNNs, especially as proxies for neurobiological computation, it is important to consider whether the resulting network structure is due to the training method itself. Here, we compare four training paradigms—BPTT, evolution strategies (ES), genetic algorithms (GA), and GA combined with Oja’s Hebbian plasticity rule (GA+Oja) — across RNNs of 32, 64, and 128 neurons. These training paradigms were applied to two contrasting tasks that are exemplary of tasks which animals perform: a discrete working memory task and a continuous sensorimotor integration task. For both tasks, we analyzed how each training algorithm modified the network through four measurements: weight change allocations across input (*W*_in_), recurrent (*W*_rec_), and output (*W*_out_) layers, the effective rank of the recurrent weight matrix, the dimensionality of hidden-state dynamics, and task accuracy. BPTT achieved near-perfect accuracy across all task conditions, progressively allocated more weight changes to *W*_out_ as the working memory task’s difficulty increased, and confined hidden-state activity to a lower-dimensional subspace than any evolutionary method. Evolutionary methods maintained higher recurrent effective rank, higher activity dimensionality, and no comparable difficulty-dependent reallocation toward the readout, all while maintaining comparable task accuracy as BPTT. These findings show how gradient-based and gradient-free algorithms discover distinct structural and dynamical solutions to the same computational problems, especially in tasks involving working memory, with important implications for analyzing task-trained RNNs as models of biological neural computation. Biological neural circuits, which are shaped by evolution and local plasticity rather than gradient descent, may operate in higher-dimensional regimes than gradient-trained RNN models predict.

## 1 Introduction

### 1.1 Recurrent Neural Networks and Biological Realism

Artificial neural networks optimized for complex ecological tasks can develop representations which closely correspond to the structural and functional coding of biological brains (Cichy et al., 2016; Yamins et al., 2014). Once these models capture these key biological features, they can serve as valuable proxies for studying real neural computation. Recurrent neural networks (RNNs) feature recurrent connections that feed the hidden state from one timestep back into the network at the next step. This looping structure enables the network to retain information from previous inputs, giving it a form of short-term memory and making it a more favored model of temporal neural computations than a purely feedforward network (Yin et al., 2021; Mienye et al., 2024). RNNs have become a dominant modeling framework in computational neuroscience, used to study how neural circuits implement cognitive functions such as working memory, decision-making, and motor control (Barak 2017; Yang & Wang 2020). Researchers train RNNs on cognitive tasks and then analyze the resulting networks to infer principles of neural computation. Trained RNNs exhibit low-dimensional recurrent dynamics during working memory maintenance, fixed-point attractors during decision-making, and oscillatory dynamics during motor control (Barak & Tsodyks 2014; Cueva et al. 2020; Ghazizadeh & Ching 2021; Mante et al. 2013; Sussillo & Barak 2013). Along with repeated demonstrations of low-dimensional recurrent dynamics in real prefrontal and motor cortical recordings at multiple scales during working memory and motor tasks (Abbaspourazad et al. 2021; Aghagolzadeh & Truccolo 2015; Gao & Ganguli 2015; Gallego et al. 2017; Inagaki et al. 2017; Li et al. 2016; Michaels et al. 2016; Sakelliadou et al. 2026; Xing et al. 2019), these findings have led to influential theories about how biological neural computations yield behavioral outputs–namely, through low-dimensional, coordinated population activity.

### 1.2 The Training Problem: Backpropagation Through Time

The prevailing methods for training RNNs are gradient-based, including backpropagation through time (BPTT), which unrolls the network across all timesteps and applies the chain rule to compute exact error gradients for every weight at every timestep (Werbos, 1990). This precise temporal credit assignment makes BPTT highly effective: it converges rapidly and achieves high accuracy on a wide range of tasks. However, BPTT faces well-known challenges. Gradients can vanish or explode as they propagate backward through many timesteps, causing instability in deep or long-sequence architectures (LeCun et al., 2015). Some workarounds exist within the gradient-based framework, such as Hessian-free optimization, which has been used to train and study RNNs as models of cortex (Martens & Sutskever 2011; Michaels et al. 2020).

More fundamentally, gradient-based methods are biologically implausible. There is no known biological mechanism that could implement the symmetric weight transport, precise error backpropagation, and global loss computation BPTT requires. Instead, the brain’s connectivity emerges from evolutionary selection operating over millions of years, combined with local synaptic plasticity rules that act within individual lifetimes (Zador, 2019). This raises a critical question that motivates the present work: if the training algorithm itself shapes the structural properties of the resulting network, how much of what we observe in gradient-trained RNNs reflects the computational requirements of the task versus the inductive biases of the optimization method?

### 1.3 Evolutionary Algorithms as Gradient-Free Alternatives

Evolutionary algorithms (EAs) provide a fundamentally different approach to training neural networks. As population-based, gradient-free search strategies inspired by Darwinian evolution, EAs maintain a population of candidate networks, evaluate their fitness on a task, and select the better performers to reproduce, accumulating useful traits over generations. This iterative cycle of selection, mutation, and reproduction enables the discovery of effective network parameters without requiring gradient information (Vikhar, 2016). EAs are particularly well-suited for optimizing neural systems where gradients are unavailable or unreliable, and early studies have demonstrated their utility as global optimizers for complex, non-differentiable problems (Slowik & Kwasnicka, 2020; Haritha et al., 2023).

Genetic algorithms (GAs) represent one of the earliest and most widely applied evolutionary approaches for neural optimization. In GAs, neural network parameters are assembled into a structured array (known as a chromosome) where each gene corresponds to a specific parameter, and the resulting population evolves through selection, crossover, and mutation. Bansal et al. (2022) demonstrated that a greedy genetic algorithm with adaptive mutation, concentrating mutation on the most impactful genes, could accelerate convergence for multilayer perceptrons. However, their approach, like most GA methods for neural networks, was designed for smaller feedforward architectures; as the number of parameters increases, the search space expands exponentially, making pure GAs impractical for large recurrent networks.

Evolution strategies (ES) occupy a middle ground between pure gradient-free search and gradient-based optimization. As formalized by Salimans et al. (2017), ES estimates the gradient from population statistics: it perturbs the current parameter vector in many random directions, evaluates each perturbation, and computes a weighted average of the perturbation directions. This produces a gradient estimate that is mathematically related to the true gradient but much noisier, especially as the number of parameters grows. Natural Evolution Strategies (NES) extend this approach by using the natural gradient to update the parameters of the sampling distribution, enabling more efficient search in high-dimensional spaces.

### 1.4 Combining Evolution with Synaptic Plasticity

Biologically plausible learning rules such as Hebbian plasticity and STDP support learning without explicit gradients but typically lack global error feedback for complex task optimization (Konishi et al., 2023). Oja’s rule, a normalized variant of Hebbian learning (Oja, 1982), maintains bounded weight growth while performing principal component extraction on neural activity patterns. The combination of evolutionary search over initial connectivity with within-lifetime Hebbian plasticity mirrors the biological separation between evolution (shaping initial connectivity over generations) and experience-dependent plasticity (adapting connectivity during an individual’s lifetime). This dual timescale of adaptation—slow evolutionary sculpting and fast synaptic tuning—is increasingly recognized as essential for understanding how biological neural circuits develop their computational properties (Zador, 2019; Kashyap et al., 2025).

### 1.5 Neuroevolution and Network Structure

A growing body of evidence suggests that gradient-free neuroevolution produces networks with qualitatively different structural properties compared to gradient-based training. Hintze and Adami (2025) demonstrated that neuroevolution tends to produce recurrent neural networks with sparser connectivity and more focused, robust information flow compared to networks trained via backpropagation. Specifically, they found that GA-optimized recurrent networks encode information in larger, more redundant groups of neurons, increasing robustness to perturbations while maintaining high information density. This contrasts with gradient descent-optimized networks, which exhibited denser but less robust connectivity patterns. Whereas Hintze and Adami (2025) characterized information flow, we quantify where the algorithm concentrates change in the network while holding task, architecture, and initialization seeds constant across four training paradigms.

These paradigm-dependent differences have important implications. If different training algorithms produce networks with different connectivity structures and dynamics but similar task outputs, then structural and dynamical features of trained networks cannot be straightforwardly interpreted as evidence about biological circuit organization. The specific structure of trained networks would be, at least in part, a signature of the training algorithm rather than a necessary feature of the computation. This possibility motivates the central question of the present work.

### 1.6 The Present Study

Earlier work has established that different optimization methods can produce qualitatively different properties in trained networks. Evolutionary algorithms offer one biologically grounded alternative to gradient-based training. Hybrid approaches combining evolutionary search with local plasticity rules are particularly promising because they approximate the multiscale learning processes (both within-lifetime synaptic plasticity and population-level natural selection) driving biological computation. Incorporating both components into a training method should support both global and local learning simultaneously and yield networks with different connectivity structures than those either method produces alone. However, a systematic comparison of how different training paradigms bias the internal connectivity structure of recurrent networks (holding architecture, task, and evaluation metrics constant) has not been conducted. Most prior work has compared methods only on task performance or only examined structural properties within a single training method.

We address this gap by comparing four training paradigms that span the gradient-information spectrum: BPTT (full gradient), ES (estimated gradient), GA (no gradient), and GA combined with Oja’s Hebbian plasticity rule (GA+Oja, combining evolutionary search with within-lifetime local plasticity). All four methods train identical RNN architectures on the same tasks to support direct comparison of the resulting connectivity structures.

Rather than focusing solely on task performance, we analyze how each training algorithm reshapes network connectivity using three complementary methods. First, per-layer weight change fractions measure the proportion of total weight change allocated to each layer (input, recurrent, output), revealing whether the algorithm preferentially modifies certain layers and whether this allocation strategy changes with task difficulty. Second, the effective rank of the recurrent weight matrix quantifies the dimensionality of the learned weight structure. Finally, principal component analysis (PCA) of recurrent population activity trajectories assesses whether structural differences in *W*_rec_ correspond to differences in dynamic dimensionality during task execution.

We evaluate all methods on two tasks: a letter n-back working memory task that scales in difficulty with the number of steps back (levels 1–4), and a two-joint arm endpoint-prediction task that requires continuous sensorimotor computation. The two tasks span two regimes of biological temporal computation: discrete working memory (n-back) and continuous sensory integration (two-joint arm), to support testing of whether connectivity signatures are task-general or task-specific. We tested network sizes from 32 to 128 neurons to examine how connectivity signatures scale with network capacity, and repeated each experiment across ten random network seeds.

We find three structural and dynamical signatures that distinguish the training paradigms despite comparable end performance. On the working memory task, BPTT progressively shifts weight changes from the recurrent to the output layer as the recall delay increases, whereas EA methods show no such difficulty-dependent reallocation. BPTT also produces recurrent weight matrices of substantially lower effective rank than any EA method, despite matched accuracy, indicating that low-rank structure is sufficient but not necessary for solving such tasks, and that selection for low-rank structure can depend on the choice of optimizer. Finally, this low-rank weight structure predicts low-dimensional population activity, while EA-trained networks occupy higher-dimensional subspaces. On the temporal integration task, the training methods converge towards more similar solutions: weight changes predominantly remain in the recurrent layer. Thus, method-dependent differences can be more or less pronounced depending on the type of task. Together, our results show that gradient-based and gradient-free training can reach the same behavior through different structural and dynamical solutions, with direct consequences for interpreting trained RNNs as models of biological computation.

## 2 Methods

### 2.1 Network Architecture

All experiments used recurrent neural networks with tanh continuous activation functions. The network contained three weight matrices: an input weight matrix *W*_in_ of dimension (*N* × obs_dim), a recurrent weight matrix *W*_rec_ of dimension (*N* × *N*), and an output weight matrix *W*_out_ of dimension (action_dim×*N*), where *N* is the number of recurrent neurons, obs_dim is the input dimension at each time point, and action_dim is the number of output dimensions at each time point. These are the exact variable names as they appear in the code. The hidden state *h*(*t*) is a vector of *N* activations that evolves according to

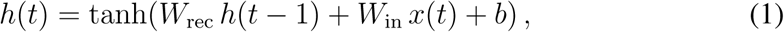

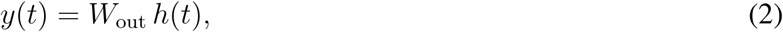

where *x*(*t*) is the input at timestep *t* and *b* is a bias vector initialized to zero; no output bias was used. The hidden state was initialized to zero at the start of each trial.

**Figure 1:**
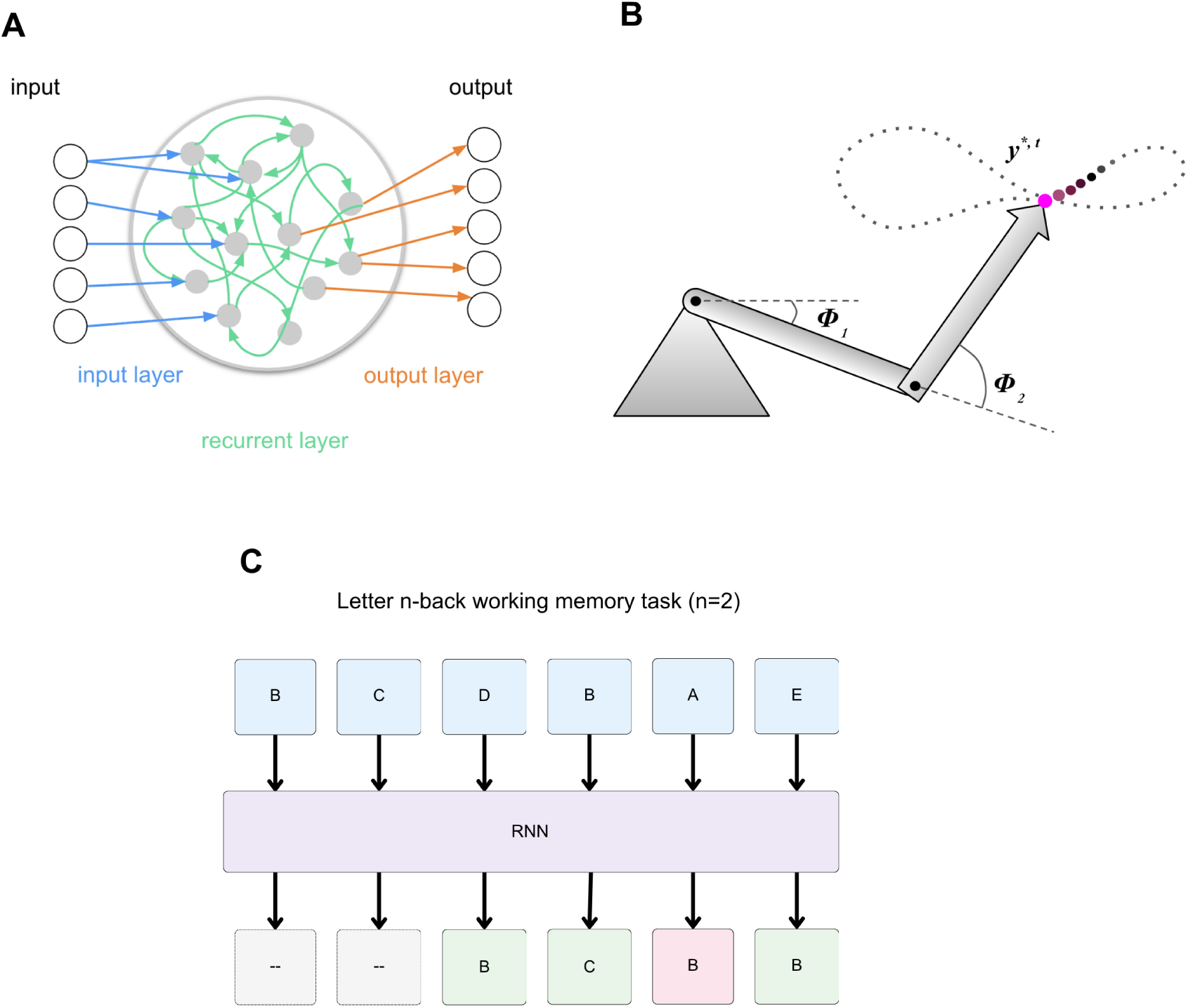
**A**. Rate-coded RNN architecture used across all four training methods. For the n-back task, five input units project to N recurrent neurons. Five output units readout from the recurrent layer. All weight matrices (W_in_, W_rec_, W_out_) are jointly trained, with bias included only in the recurrent layer. **B.** Two-joint planar arm task. The arm has two unit-length links with joint angles φ_1_ and φ_2_, integrated from angular velocity inputs ω_0_ and ω_1_. The magenta dot marks the endpoint position y*(t), which traces a smooth, nonlinear trajectory (dotted) over 20 timesteps. At each timestep, the network receives the two angular velocity inputs and must predict the (x, y) endpoint position using forward kinematics. **C.** Schematic of the letter n-back task at difficulty level n=2. At each timestep, the network receives an input letter (A–E) and must output the letter 2 steps prior in time. Input letter_1_s_1_(in blue) are presented sequentially to the RNN, which updates its hidden state at each timestep. Output responses of the RNN

We tested network sizes of *N* = 32, 64, and 128 neurons. For a network of *N* neurons with obs_dim input dimensions and action_dim output dimensions, the total number of trainable parameters is *N* ^2^ + *N* ×obs_dim+action_dim× *N* + *N* (including bias). For the n-back task (obs_dim = action_dim = 5), this yields 1,184 parameters at *N* = 32, 4,480 at *N* = 64, and 17,408 at *N* = 128. The quadratic growth of the recurrent weight matrix dominates this scaling and is the central reason evolutionary methods require dimension-aware hyperparameter scaling at larger N.

All weight matrices were initialized from a normal distribution with standard deviation 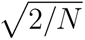. The same random initialization was used across all four training methods for each seed (refer to Experimental Design and Replication), so any differences in the final trained networks are attributable to the training algorithm rather than differences in starting conditions.

For BPTT, the identical architecture was implemented in PyTorch to enable automatic differentiation through the computational graph. For ES, GA, and GA+Oja, the architecture was implemented in NumPy with explicit forward-pass computation, since these methods do not require gradient computation.

### 2.2 Letter N-back Task

The letter n-back recall task requires the network to recall an input from n timesteps ago within a 20-timestep sequence, maintaining a working memory of preceding inputs. At each timestep in a trial, one of five symbols (A, B, C, D, E) was presented as a one-hot vector of dimension 5. The network produced a 5-dimensional vector as its prediction of the symbol that appeared n timesteps earlier; this output is passed through a softmax function to generate a probability distribution across the five symbols. The loss is the cross-entropy between the one-hot target vector and the probability vector and averaged over the 20 timesteps. Accuracy is the percentage of the timesteps where the network’s argmax output matches the target symbol. For the first n timesteps of each trial (before there exists a valid recall target n steps back), the target is set to the current input, providing a mapping that does not require memory. We tested n=1 through n=4; difficulty increases with n because the network must maintain the symbol over a longer delay.

The choice of output encoding was critical for ensuring fair comparison across methods. For example, one could use a single scalar output, encoding the target symbol as a number from 0 to 4, which will allow BPTT to achieve high accuracy by primarily tuning *W*_out_ (learning a linear mapping from the hidden state to a scalar) without meaningfully restructuring recurrent dynamics. The five-unit one-hot encoding with cross-entropy loss prevents this shortcut: the network must compute and maintain a distributed representation of the letter identity in its hidden state, requiring genuine recurrent computation.

Each training method evaluated fitness on 20 randomly generated trials per update step. For evolutionary methods (ES, GA, GA+Oja), one evaluation (see *Fitness Evaluation and Forward Pass*) consists of a single forward pass of one candidate weight vector through all 20 trials to compute its fitness score; this is repeated for each individual in the population before any weight update occurs. The equivalent process of an evaluation in BPTT is one iteration, in which a forward pass over a batch of 20 trials occurs, followed by a single Adam gradient step.

Symbols were drawn uniformly at random with replacement at each timestep. Within each evaluation, the same random seed controlled trial generation to reduce noise in fitness comparisons across candidates, but different trials were generated across evaluations (for evolutionary methods) or iterations (for BPTT) to prevent overfitting to specific sequences.

### 2.3 Two-joint Arm Endpoint Prediction Task

The two-joint arm task uses a two-link planar arm model where each link has unit length (Scherr et al. 2020). Two joint angular velocities (*ω*_0_, *ω*_1_) serve as input at each timestep, and the network must predict the (*x, y*) position of the arm’s endpoint. The angular velocities are generated as sums of five sinusoids with random frequencies, phases, and amplitudes, producing smooth but complex trajectories. Joint angles are computed by cumulatively integrating the angular velocities over time, *φ*(*t*) = Δ*t* Σ *ω*(*t^′^*), and the endpoint position is derived from forward kinematics:

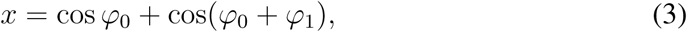

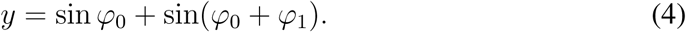

Both inputs and targets are normalized to [−1, 1].

The loss function was mean squared error (MSE) between the network’s output and the target endpoint position, averaged over all 20 timesteps. To report accuracy on a comparable scale to the n-back task, we convert MSE to a percentage using exp(−MSE), which ranges from 0% (high error) to 100% (zero error). Unlike the n-back task, this task requires continuous regression rather than discrete classification, and involves tightly coupled dynamics: the output at each timestep depends on the integral of all previous inputs rather than a single stored symbol. This tests whether connectivity signatures that emerge in trained networks are shared across qualitatively different computational domains.

Sequence length was 20 timesteps, matching the n-back task. Each trial generated a new random trajectory with different sinusoidal parameters. Fitness was evaluated over 20 trials per evaluation, identical to the n-back protocol.

### 2.4 Training Procedure: Backpropagation Through Time (BPTT)

The BPTT pipeline trained a single network using gradient-based optimization. The network was implemented as a PyTorch *nn.Module* with *W*_rec_, *W*_in_, and *W*_out_ as *nn.Parameter* objects, enabling automatic differentiation through the entire computational graph. The hidden state was initialized to zero at the start of each trial. On each training iteration, a batch of 64 trials was generated, each consisting of a 20-timestep input sequence and the corresponding target sequence. Gradients are accumulated over the full 64-trial batch into one scalar loss, then a single optimizer.step() updates all weights once per iteration, so nothing carries across iterations.

The forward pass unrolled the recurrent computation explicitly across all 20 timesteps. At each timestep *t*, the hidden state was updated as 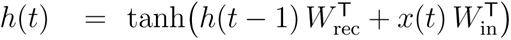, and the output was computed as 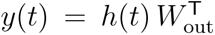, producing raw logits of dimension action_dim. For the n-back task, these logits were passed through a softmax and compared to the one-hot target with cross-entropy loss, averaged over all 20 timesteps and all 64 trials in the batch. For the two-joint arm task, the two-dimensional output was compared directly to the target endpoint using mean squared error; no output activation was applied.

Parameter updates used the Adam optimizer (Kingma & Ba, 2015) with learning rate 0.001, *β*_1_ = 0.9, *β*_2_ = 0.999, and *ε* = 10*^−^*^8^. No learning rate scheduling, weight decay, or dropout regularization was applied.

Training ran for 2,000 iterations at *N* = 32 and *N* = 64, and 500 iterations at *N* = 128. BPTT consistently converged within 50–200 iterations at all sizes, so the longer training at smaller sizes served only to confirm convergence stability. Accuracy was computed every 25 iterations on a fresh batch of 20 trials not used for gradient computation. The initial weights (*W*_init_) and final weights (*W*_final_) were saved for each run, along with the full training history (loss and accuracy per iteration).

### 2.5 Training Procedure: Evolution Strategy (ES)

The ES pipeline implemented the OpenAI Evolution Strategy (Salimans et al., 2017). The algorithm maintains a single parameter vector *θ* (the flattened concatenation of *W*_rec_, *W*_in_, and *W*_out_) and iteratively updates it using a gradient estimate derived from population evaluations.

**Figure 2:**
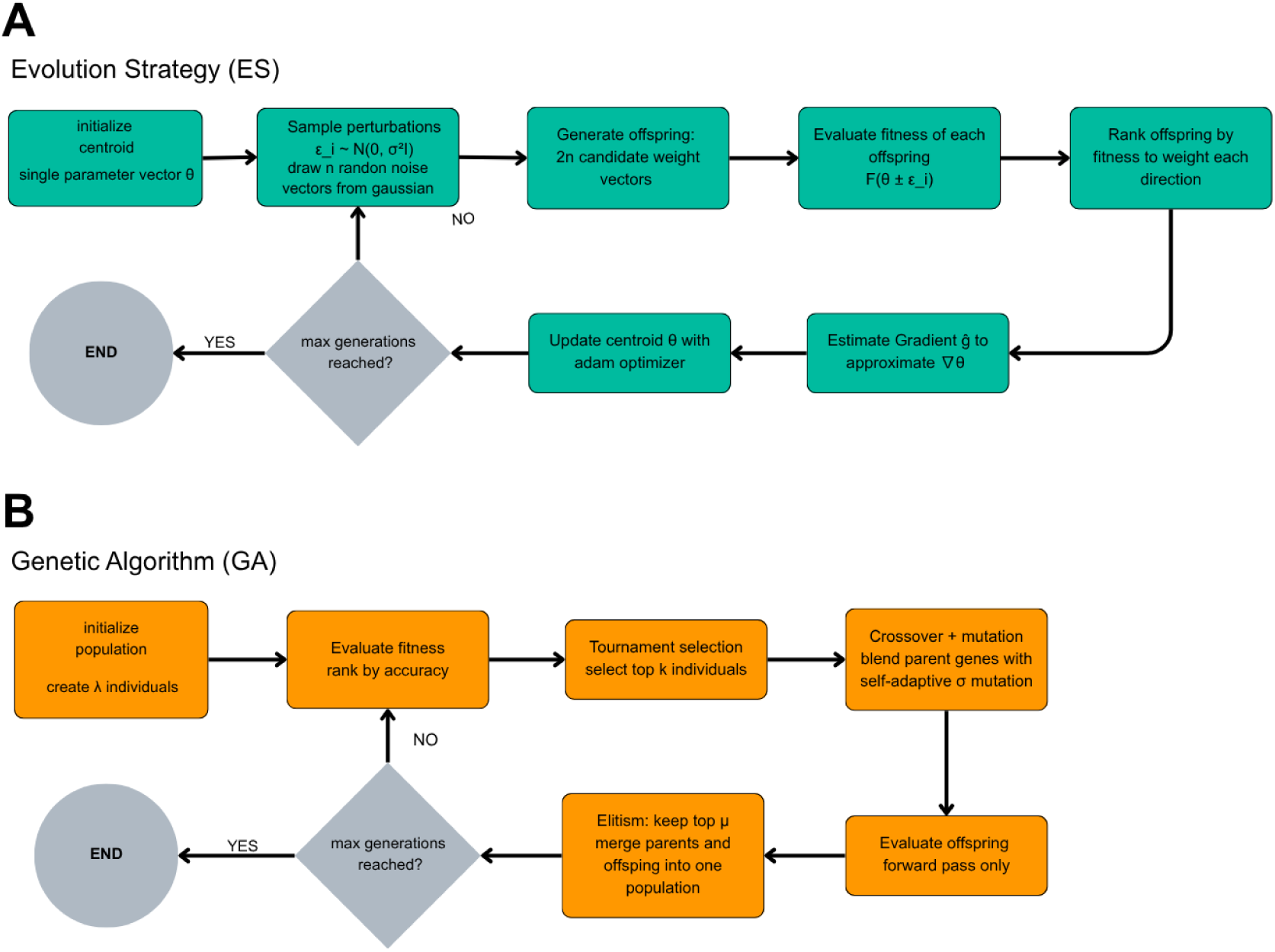
**A.** Evolution Strategy (ES) Training Procedure. Starting from a single parameter vector θ, perturbations ε_i ∼ N(0, σ^2^I) are sampled each generation to produce mirrored (antithetical) pairs. Each candidate is evaluated on the task, and a weighted average of perturbation directions estimates the gradient ∇θ. The centroid is updated via Adam optimizer and σ is adapted using Rechenberg’s 1/5-success rule. This process repeats until the generations are complete. **B.** Genetic Algorithm (GA) training procedure. A population of λ individuals is randomly initialized and evaluated by task fitness. Parents are chosen using tournament selection, whose genes are recombined using neuron blend crossover and self-adaptive σ mutation. Elitism preserves the top µ individuals unchanged in the next generation. This cycle repeats until the generation limit is reached.

At each generation, the algorithm sampled Gaussian noise vectors *ε_i_* ∼ N (0*, I*) and evaluated both *θ* + *σε_i_* and *θ* − *σε_i_*(mirrored sampling), where *σ* is the perturbation standard deviation. Each perturbed parameter vector was decoded into weight matrices, a network was instantiated with those weights, and fitness was evaluated by averaging the negative loss across 20 random trials. The gradient estimate was then computed, and the parameter vector updated, as

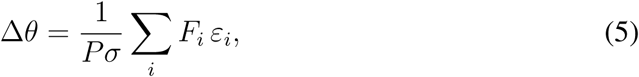

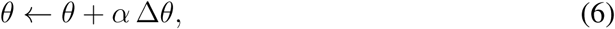

where *F_i_*is the fitness of the *i*-th perturbation, *P* is the population size, and *α* is the learning rate. Population size and *σ* were scaled with network size to preserve comparable search dynamics across sizes, as detailed below.

The perturbation standard deviation *σ* was adapted during training using Rechenberg’s 1/5-success rule: the algorithm tracked whether the best fitness improved over a rolling window of 20 generations. If the success rate exceeded 1/5, *σ* was increased by a factor of 1.10 to encourage broader exploration; if it fell below 1/5, *σ* was decreased by a factor of 0.95 to focus the search. *σ* was bounded between 0.005 (or 25% of the initial value, whichever was larger) and 4× the initial value.

The initial *σ* value of 0.02 was determined by a preliminary hyperparameter sweep at *N* = 32 on 1-back that varied *σ* (0.005, 0.01, 0.02, 0.05, 0.1, 0.2), population size (128, 256), and learning rate (0.01, 0.03, 0.05, 0.1). The sweep revealed that *σ* was the most critical variable: the default value of 0.1 was too large for effective gradient estimation, with perturbations overwhelming the signal. The best *N* = 32 configuration (*σ* = 0.02, population = 256, learning rate = 0.03) achieved 65.5% accuracy on 1-back in this preliminary sweep.

For the main experiments, ES used a learning rate of 0.03 and ran for 500 generations, with both *σ* and population size scaled with network size to preserve search dynamics across the quadratically growing parameter space. Population size was set to 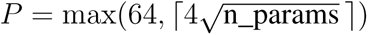, yielding *P* = 147 at *N* = 32, *P* = 268 at *N* = 64, and *P* = 528 at *N* = 128. The perturbation scale was set to 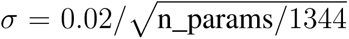, where 1344 is the *N* = 32 parameter count for the n-back task, yielding *σ* = 0.02 at *N* = 32, *σ* = 0.011 at *N* = 64, and *σ* = 0.006 at *N* = 128. Without these scalings, ES suffered performance collapse at larger N, which is attributable to fixed hyperparameters and not necessarily a fundamental limitation of the algorithm.

The initial parameter vector served as *W*_init_ for connectivity analysis. The final parameter vector after 500 generations was decoded into 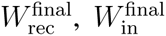, and 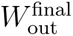. The full history of mean and best fitness, mean and best accuracy, and *σ* values per generation was saved.

### 2.6 Training Procedure: Genetic Algorithm (GA)

The GA pipeline implemented an elitist genetic algorithm with a (*µ* + *λ*) selection scheme, in which *µ* = 4 elites survived each generation and were augmented by *λ* = *P* − 4 offspring, where *P* is the total population size (see below). The algorithm combined tournament selection, neuron-level blend crossover, and self-adaptive mutation. Each individual was represented as a genotype with two parts: a weight gene vector containing the flattened *W*_rec_, *W*_in_, and *W*_out_ matrices (totaling *N* ^2^ + *N* × obs_dim + action_dim × *N* entries), and a *σ* gene vector of *N* per-neuron mutation-rate parameters.

Population size *P* was scaled with the parameter count to preserve search dynamics across network sizes: 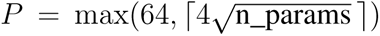, giving *P* = 147 at *N* = 32, *P* = 268 at *N* = 64, and *P* = 528 at *N* = 128. Weight genes were sampled from a normal distribution with standard deviation 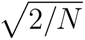, matching the BPTT initialization; *σ* genes were all initialized to 0.05. The population centroid (the element-wise mean of all individuals’ weight genes) was saved as *W*_init_ for connectivity analysis, providing a reference point comparable to BPTT’s single-model initialization.

Each generation proceeded through five steps. First, all individuals were evaluated: each genotype was decoded into weight matrices, instantiated as a network, evaluated on 20 random trials, and assigned a fitness equal to the mean negative loss across trials. Second, fitness sharing was applied: each individual’s raw fitness was divided by a niche count (the number of other individuals within a genotypic distance threshold), penalizing crowded regions of the search space to maintain diversity. Third, parents were selected by tournament selection with tournament size *k* = 5: five individuals were sampled uniformly at random from the population, and the one with the highest shared fitness was chosen. Fourth, *λ* = *P* −4 offspring were generated by crossover and mutation. Fifth, the top 4 individuals by **raw** fitness (not shared fitness) were preserved unchanged as the next generation’s elites.

Crossover used a neuron-level blend mechanism. For each of the *N* neurons, a blend coefficient *α_i_* was sampled uniformly from [0.3, 0.7]. The *i*-th row of the offspring’s *W*_rec_ was set to *α_i_* (parent1 row) + (1 − *α_i_*) (parent2 row), and similarly for the *i*-th row of *W*_in_ and the *i*-th column of *W*_out_. This preserves the neuron as a functional unit during recombination, which is more biologically motivated than gene-level crossover, which would scramble the relationships between a neuron’s incoming and outgoing connections. *σ* genes were blended using the same per-neuron coefficients.

Mutation was self-adaptive at two levels. At the individual level, each neuron carried a per-neuron mutation-rate gene *σ_i_* that evolved alongside the weight genes it controlled. At each mutation event, *σ_i_* was first updated via a lognormal perturbation, *σ_i_* ← *σ_i_*exp(*τz*), where *z* ∼ N (0, 1) and 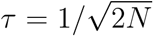 is the Schwefel self-adaptation learning rate. The *σ_i_* values were clamped to [0.005, 0.15] to prevent mutation rates from vanishing (stagnation) or exploding (random search). The effective mutation rate for neuron *i* was then scaled by a rank-based multiplier based on an individual’s raw-fitness rank: the top quartile received 0.5× (fine-tuning near the best solutions), the middle half received 1.0×, and the bottom quartile received 2.0× (broader exploration for poor performers). For each weight associated with neuron *i*, mutation was applied stochastically: if a uniform random draw fell below the effective *σ_i_*, the weight was perturbed by adding Gaussian noise scaled by the global mutation standard deviation.

At the population level, the global mutation standard deviation was adapted with Rechenberg’s 1/5-success rule, analogous to the ES *σ* adaptation. Over a rolling 20-generation window, if the best fitness improved in more than 20% of generations the global mutation standard deviation was increased by a factor of 1.05; if it improved in fewer than 20%, it was decreased by a factor of 0.97. The amplitude was bounded between 25% and 400% of its initial value. To keep per-weight perturbations proportional to the initialization scale across network sizes, the initial global mutation standard deviation was scaled as 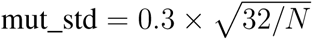, giving 0.30 at *N* = 32, 0.21 at *N* = 64, and 0.15 at *N* = 128. Without this scaling, the same nominal amplitude of 0.3 would be ∼2.4× the initialization scale at *N* = 128 versus ∼1.2× at *N* = 32, making mutations disproportionately disruptive at larger sizes.

Each generation produced *P* −4 offspring to fill the population alongside the 4 elites. The GA ran for 500 generations at all network sizes. The best individual’s weight genes were then decoded into 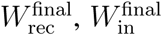, and 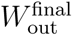. The full training history (mean and best fitness, mean and best accuracy, and mean *σ* per generation) was saved.

### 2.7 Training Procedure: GA with Oja’s Hebbian Plasticity (GA+Oja)

**Figure 3:**
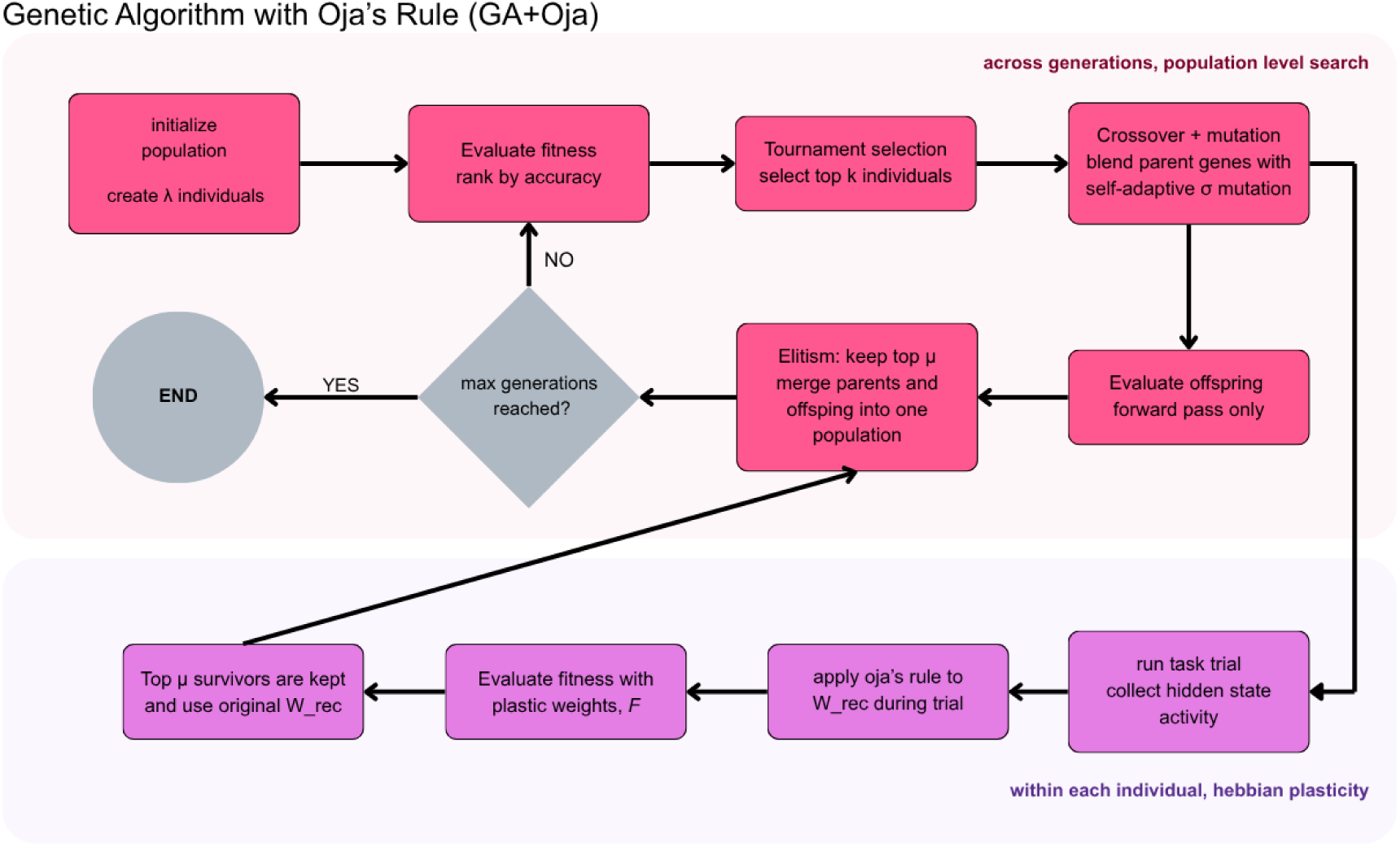
GA with Oja’s Hebbian plasticity (GA+Oja) training procedure, illustrating the two-timescale scheme. GA standard population-level search (pink) evolves the initial weights and plasticity parameters η and w_max_. Oja’s rule (purple, within each individual’s evaluation) modifies a per-trial copy of W_rec_ at every timestep using the current hidden-state activity. Fitness is computed using Oja’s evolved weights. The original genotype W_rec_ is restored after each Oja’s trial with no Lamarckian inheritance.

The GA+Oja pipeline used identical GA mechanics to those described above (same population size, selection, crossover, mutation, elitism, and fitness sharing), with one critical addition: each fitness evaluation incorporated within-trial Hebbian plasticity on the recurrent weight matrix via Oja’s rule.

The genotype extended the standard GA format with two additional genes per individual: log *η* (the log-transformed Hebbian learning rate) and log *w*_max_ (the log-transformed weight clipping bound). These decoded to *η* = exp(log *η*), clipped to [10*^−^*^6^, 1.0], and *w*_max_ = exp(log *w*_max_), clipped to [0.1, 20.0]. The Oja parameters were mutated in log-space with the same global mutation standard deviation used for weight genes, and were blended during crossover using a single global blend coefficient sampled uniformly from [0.3, 0.7].

During each fitness evaluation, the individual’s weight genes were decoded into weight matrices and a working copy of *W*_rec_ was made; this copy (not the genotype’s *W*_rec_) was modified by Oja’s rule during the trial. At each of the 20 timesteps, once the hidden state *h*(*t*) was computed from the current *W*_rec_ copy, the Oja update was applied as

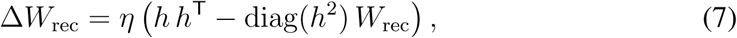

where *h* is the *N* -dimensional activation vector. The updated *W*_rec_ was then clipped element-wise to [−*w*_max_*, w*_max_]. This cycle repeated at every timestep of the trial.

Critically, the Oja-modified *W*_rec_ was used only for computing the network’s output and thus its fitness score during that trial. After the trial ended, the modified *W*_rec_ was discarded; the genotype’s original *W*_rec_ was preserved unchanged for the next generation. This means there was no Lamarckian inheritance: within-trial plasticity changes did not feed back into the evolutionary process. Evolution operated on the initial weights and the plasticity parameters, while Oja’s rule provided within-trial adaptation. This design mirrors the biological separation between evolutionary shaping of initial connectivity (across generations) and experience-dependent plasticity (within a lifetime).

### 2.8 Fitness Evaluation and Forward Pass

All four training methods used the same loss / fitness metric for each task. For the n-back task, fitness was defined as the negative cross-entropy loss averaged across all timesteps and all evaluation trials; for the two-joint arm task, it was defined as the negative MSE:

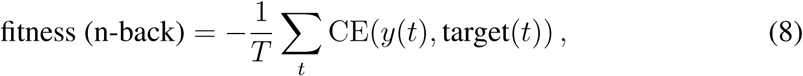

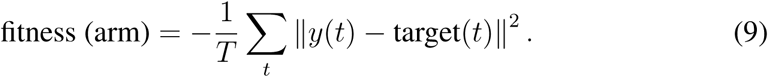

Higher fitness (closer to zero) indicates better performance. Each evaluation consisted of 20 randomly generated trials. For BPTT, the batch size was 64 trials per iteration, but accuracy was evaluated on 20 fresh trials every 25 iterations for comparability with the evolutionary methods.

For each trial evaluation (across all methods), the network’s forward pass proceeded identically. The hidden state *h* was initialized to zero. At each timestep *t* = 0, 1, …, 19, the input *x*(*t*) was a one-hot vector of dimension 5 (n-back) or a 2-dimensional angular velocity vector (two-joint arm). The hidden state was updated as *h*(*t*) = tanh(*W*_rec_ *h*(*t* − 1) + *W*_in_ *x*(*t*)), and the output was computed as *y*(*t*) = *W*_out_ *h*(*t*). For the n-back task, y(t) was a 5-dimensional logit vector; during loss computation, softmax was applied to convert to probabilities, and cross-entropy was computed against the target class. For the two-joint arm task, y(t) was a 2-dimensional (x, y) prediction and MSE was computed directly. In GA+Oja, the *W*_rec_ used in this forward pass was the Oja-modified copy (updated at each timestep), while *W*_in_ and *W*_out_ remained fixed throughout the trial.

For the n-back task, accuracy was the fraction of timesteps on which the network’s argmax output matched the target class, excluding the first n warm-up timesteps (before a valid n-back target exists). For the two-joint arm task, accuracy was defined as exp(−MSE), yielding a percentage-scale metric where 100% corresponds to zero error and lower values indicate higher prediction error.

### 2.9 Connectivity Analysis

For each trained network, connectivity analysis proceeded as follows. The weight change for each layer was computed as 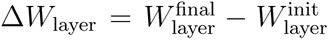. Per-layer weight change fractions were computed as 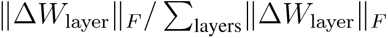, where 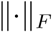 denotes the Frobenius norm (the square root of the sum of squared elements). This metric captures the relative allocation of learning effort across network layers, independent of the total magnitude of weight change. Because we divide each layer’s change by the sum of changes across layers, two networks that changed by very different amounts (learning quickly with large weight updates vs. slow convergence with small updates) will have the same fractions if they allocated proportionally the same effort to each layer. This method allows for direct comparison of where each algorithm concentrated its learning across methods and conditions without conflating allocation strategy with overall learning magnitude. For BPTT, *W*_init_ and *W*_final_ are the single model’s weights before and after training. For ES, *W*_init_ is the initial parameter vector and *W*_final_ is the final parameter vector after 500 generations. For GA and GA+Oja, *W*_init_ is the population centroid at generation 0 and *W*_final_ is the best individual’s weights at the end of evolution.

Effective rank of the recurrent weight matrix was computed from its singular value decomposition (SVD). Following Roy and Bhatt (2007), effective rank is defined as

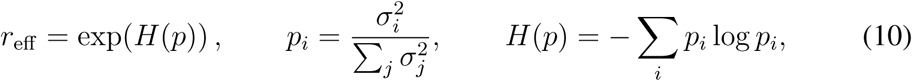

where *p_i_* is the proportion of total squared variance explained by the *i*-th singular value and *H*(*p*) is the Shannon entropy of this distribution. A matrix with all singular values equal attains the maximum effective rank equal to its dimension, indicating that recurrent connectivity spans the full space; a matrix dominated by a few large singular values has low effective rank, indicating that recurrent connectivity is confined to a low-dimensional subspace. For a 32-neuron network, the theoretical maximum effective rank is 32. Across 10 random seeds, untrained *W*_rec_ (normal initialization, 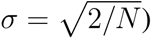 has an effective rank of 16.6 ± 0.5 at *N* = 32, 32.9 ± 0.3 at *N* = 64, and 65.7 ± 0.5 at *N* = 128, providing a baseline against which to read the trained values.

### 2.10 Dynamical Analysis: Activity Dimensionality

To assess how the choice of training paradigm affects the dimensionality of recurrent network dynamics, we performed principal component analysis (PCA) on the recurrent (hidden) layer activity of trained networks. Trajectories were collected by running eight trials of the task through each trained model without any further training. Eight trials across 20 timesteps yield 160 pooled timepoints, exceeding the largest network size (*N* = 128) so the covariance is full-rank estimable for all networks. At each timestep within a trial, the hidden state vector *h*(*t*) ∈ ℝ*^N^* was recorded to generate an activity matrix of shape (total_timesteps × *N*). PCA decomposes the *N* × *N* covariance matrix of neural activity. Pooling across trials supports covariance in capturing fluctuations across both within-trial dynamics and across-trial input variation to record the full range of visited states rather than a single trajectory’s geometry. In a single trial, PCA focuses more on the trajectory geometry of that specific input. A random fixed seed drew trials for reproducibility and comparability across methods and network sizes.

Prior to decomposition, the activity matrix was mean-centered. PCA used singular value decomposition (SVD) of the centered matrix. The number of principal components required to explain 90% of total variance in the hidden-state trajectories was used as the metric of effective dimensionality. The compression ratio is defined as the number of PCs required for 90% explained variance divided by the network size N to allow for direct comparison across network sizes.

PCA was conducted across all four methods, four n-back difficulty levels, the two-joint arm task, and network sizes of 32, 64, and 128, with 10 network seeds for each condition. Results were aggregated across all ten random seeds; values are reported as mean ± standard deviation per condition. The analysis was implemented in scripts/analyze_activity_pca.py.

### 2.10 Experimental Design and Replication

All experiments were repeated with ten random seeds (42, 123, 456, 789, 1011, 1213, 1415, 1617, 1819, 2021). Each seed controlled both the weight initialization and the random trial generation for reproducibility. For each seed, the same initial weight matrices were used across all four training methods: the ES and BPTT pipelines initialized their single parameter vector or model directly from the seed; the GA and GA+Oja pipelines initialized their population using the seed, and the population centroid served as the comparable “initial weights” for connectivity analysis. Results are reported as mean ± standard deviation across seeds.

For each run, the following artifacts were saved to disk: the initial weight matrices 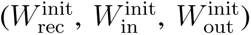, the final weight matrices 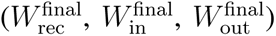, the full training history (fitness, accuracy, and method-specific diagnostics per iteration or generation), and a JSON configuration file specifying all hyperparameters. For GA+Oja, an additional 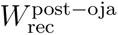 matrix was saved. Connectivity analysis figures (weight change fractions, delta distributions, recurrent weight heatmaps, final weight distributions, and sparsity comparisons) were generated automatically after each run using a shared analysis script.

The full experimental sweep comprised 480 training runs for the n-back task (4 methods × 4 n-back levels × 3 network sizes × 10 seeds) and 120 runs for the two-joint arm task (4 methods × 3 network sizes × 10 seeds). Cross-seed analysis scripts aggregated results to compute summary statistics and generate the figures presented here.

All pairwise comparisons used Mann-Whitney U tests (non-parametric; n = 10 per group), and trend analyses used Spearman rank correlations. Reported p-values are Holm–Šidák adjusted across the full family of 78 simultaneous tests. Effect sizes are reported as rank-biserial correlation (*r*_rb_) for Mann-Whitney tests and Spearman *ρ* for trend tests. Bootstrap 95% confidence intervals were computed from 2,000 resamples.

### 2.11 Software and Computational Resources

The source code, configuration files, and analysis scripts are publicly available at https://github.com/mivyer/rnn-training-paradigms. The repository includes the run_experiment.py entry point for reproducing individual runs, sweep scripts for running the full experimental matrix, and analysis scripts (ana-lyze_cross_seed.py, analyze_two-joint_t20.py, make_cns_figure.py) for reproducing all figures.

## 3 Results

### 3.1 Performance on the N-back Task

BPTT achieved near perfect accuracy (≥ 99.8%) at n-back levels 1-4 across all network sizes (Table 1). ES and GA also achieved high accuracy at 1- and 2-back across all sizes (≥ 98.0% in every condition), with performance diverging at harder levels and smaller network sizes. At *N* = 32 and 4-back, ES dropped to 58.6% ± 21.6% and GA to 82.8% ± 8.8%, reflecting the limitations of gradient-free optimization for longer-range temporal credit assignment. GA+Oja consistently underperformed plain GA at all network sizes and difficulty levels (Mann-Whitney U, *p*_adj_ < 0.001 at all three network sizes). Although EA accuracy collapses at *N* = 32/64, it recovers fully at *N* = 128, so smaller network accuracy might be a result of scaling effects.

**Table 1:**
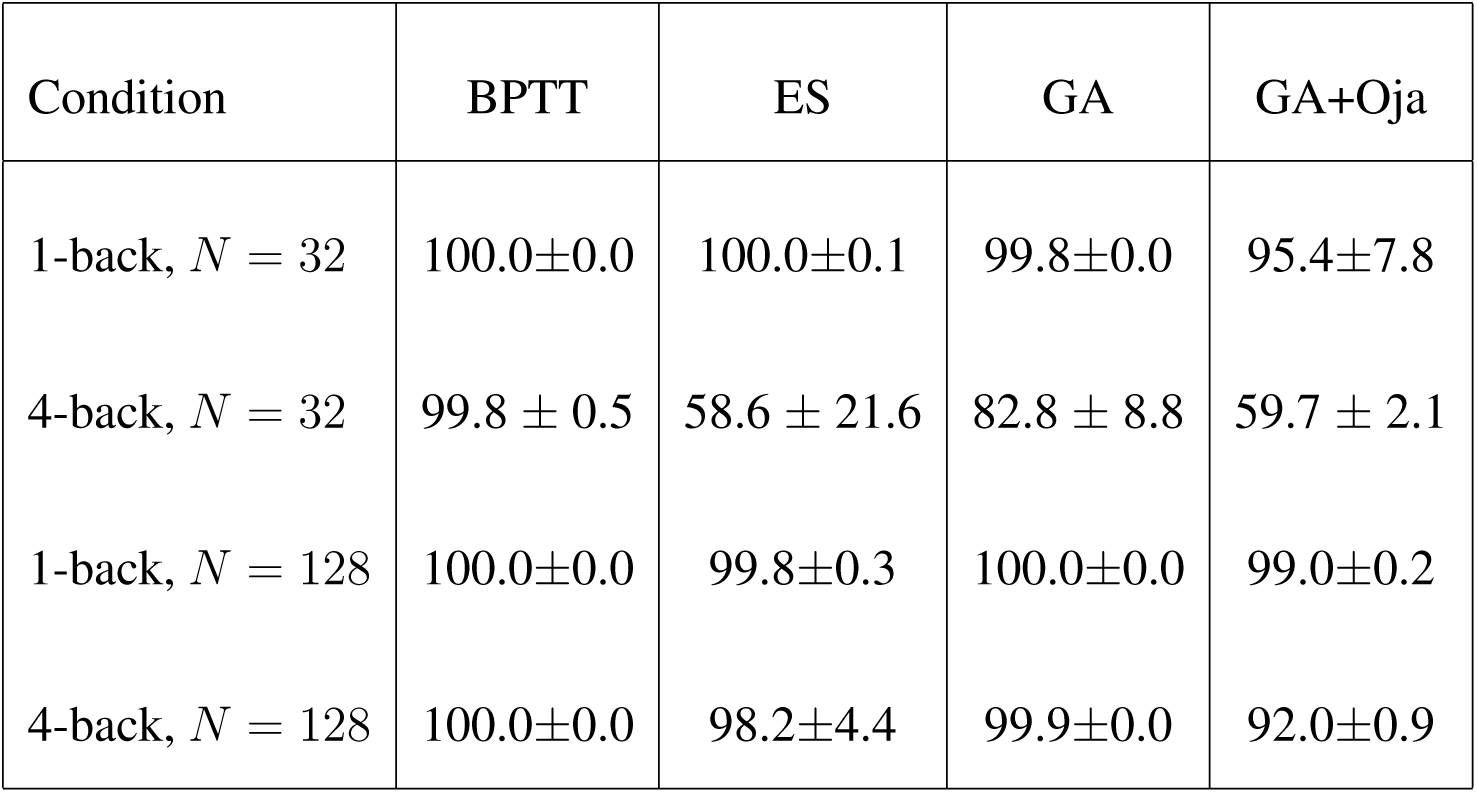
Accuracy at the easiest and hardest n-back levels for the smallest and largest network sizes tested (mean ± std across 10 seeds). The *N* = 32, 4-back row captures the performance collapse of gradient-free methods at the most difficult condition; the *N* = 128 rows show recovery under dimension-aware hyperparameter scaling.

Accuracy improved with network size for all evolutionary methods (Spearman *ρ*: ES = +0.453, p = 0.012; GA = +0.943, p < 0.001; GA+Oja = +0.736, p < 0.001), under dimension-aware hyperparameter scaling described in Methods. GA reached 99.9% ± 0.0% at *N* = 128 4-back and ES recovered to 98.2% ± 4.4%, demonstrating that the performance collapse at smaller sizes is a scaling effect rather than a fundamental limitation of gradient-free optimization. BPTT showed no sensitivity to network size, achieving near-perfect accuracy at all scales.

**Figure 4:**
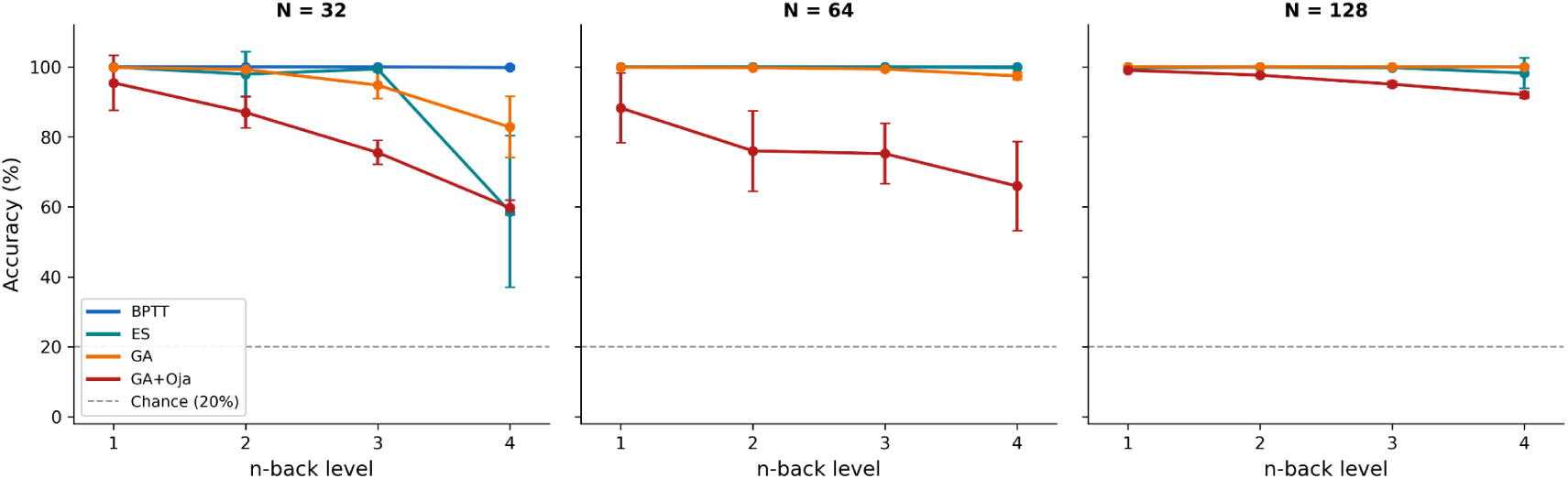
Performance vs. task difficulty on the letter n-back task (N = 32, 64, 128 neurons; 10 seeds; mean ± std). BPTT achieves ≥ 99.8% accuracy at all n-back levels and network sizes. ES collapses at N = 32, 4-back (58.6% ± 21.6%) before recovering at N = 64 and N = 128 under dimension-aware hyperparameter scaling. GA shows improvement with network size on 4-back (Spearman ρ=+0.943, p<0.001), reaching 99.9% at N = 128 4-back. GA+Oja consistently underperforms GA. Dashed line indicates chance performance (20%).

### 3.2 Effective Rank of *W*_rec_

The most striking structural difference between training paradigms was in the effective rank of the recurrent weight matrix (Fig 5). Compared to the random-initialized rank baseline of 16.6 ± 0.5 at *N* = 32 (theoretical maximum of 32), BPTT compressed effective rank far below its starting point: 8.4 at 1-back, rising to 10.2 at 4-back. All three evolutionary methods, in contrast, maintained baseline effective ranks between 15.9 and 16.6 across difficulty levels, even though they substantially modified *W*_rec_ in magnitude (Section 3.3). BPTT is the only method which shifts recurrent rank away from initialization, Every pairwise comparison between BPTT and each evolutionary method was statistically significant at every n-back level and every network size (Mann–Whitney U; *r*_rb_ = 1.00 and *p*_adj_ < 0.004 in all 36 comparisons).

**Figure 5:**
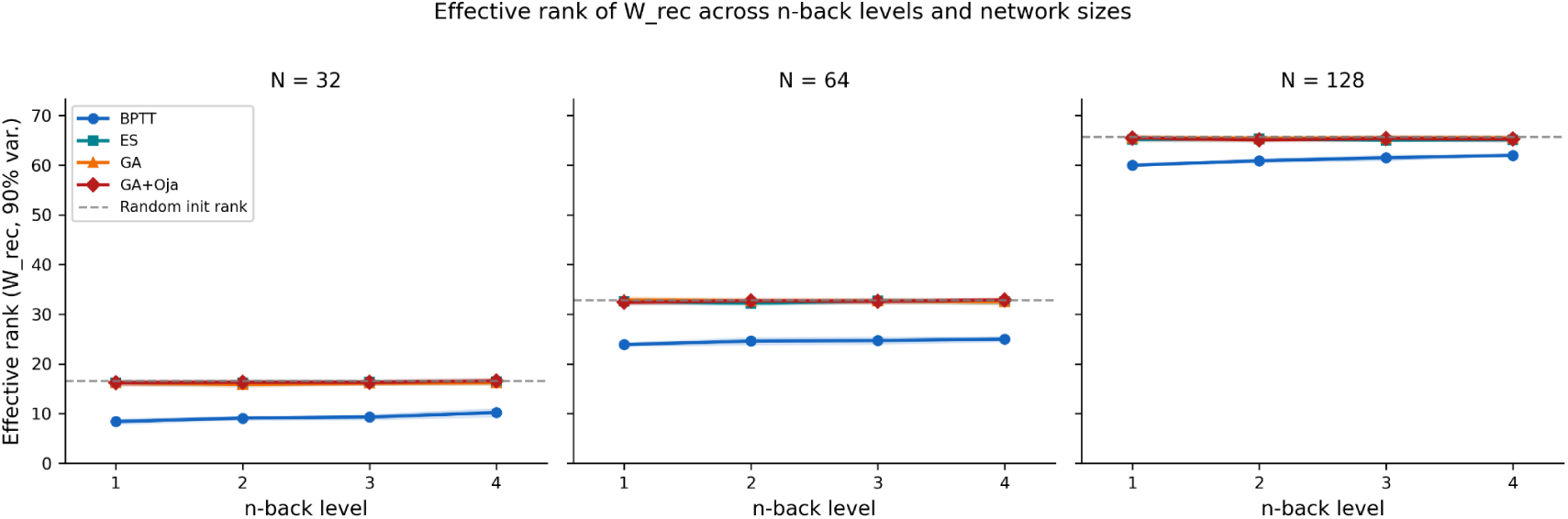
Effective rank of W_rec_ by method and n-back level. The dashed line marks the random-initialization baseline (16.6, 32.9, 65.7). Evolutionary methods track the baseline at all sizes; only BPTT falls below it, most steeply at the smallest size. Every BPTT-vs-EA comparison is significant (Mann–Whitney U, r_rb_ = 1.00, p_adj_ < 0.004 across all 36 comparisons).

At *N* = 64 (baseline 32.9 ± 0.3), BPTT produced effective ranks of 23.9-25.0, compared to EA methods clustering at 32.5-32.7. At *N* = 128, BPTT networks (baseline 65.7 ± 0.5) ranged in recurrent rank from 60.0-62.0, while EA methods stayed at 65.0-65.5. The BPTT to EA ratio narrows with increasing network size, with the ratio of BPTT to EA at approximately 0.52 at *N* = 32, 0.76 at *N* = 64, and 0.94 at *N* = 128. The ordering (BPTT < EA) and its statistical significance hold for every network size, even as the magnitude of the gap shrinks with N.

This effective-rank difference is not explained by performance differences. At 1-back and 2-back, BPTT, ES, and GA all achieve ≥ 97.9% accuracy at *N* = 32 yet still show the full rank separation, confirming that the gap reflects the training algorithm rather than differential task difficulty. The low rank of BPTT networks is consistent with the implicit bias of gradient descent toward low-rank solutions documented in the optimization theory literature (Gunasekar et al., 2017; Arora et al., 2019): gradient updates concentrate along a few principal directions in weight space. Evolutionary methods, lacking this inductive bias, retain the near-baseline rank of the naive network.

### 3.3 Per-Layer Weight Change Fractions

Consistently, the majority of weight changes occur in *W*_rec_ during training, as it is the largest layer and critical in maintaining the memory trace necessary for the task. However, changes in *W*_out_ increase with task difficulty: in BPTT-trained networks, at increasing n-back levels for *N* = 32, the fraction of total weight change allocated to *W*_out_ increased from 27.4% to 37.9% (Spearman *ρ* = +0.969, *p*_adj_ < 0.001; bootstrap 95% CIs: [0.273, 0.276] at 1-back to [0.374, 0.384] at 4-back). *W*_in_ weight changes decreased correspondingly, and *W*_rec_ weight changes remained stable at 47.8–50.5%.

This trend followed across all network sizes: *ρ* = +0.969 at *N* = 64 and *ρ* = +0.941 at *N* = 128 (both *p*_adj_ < 0.001; Fig 6). The bias towards allocating learning in the output weights is consistent with BPTT’s gradient signal concentrating learning in the layer with the most direct influence on loss. This readout-refinement strategy is almost certainly an attribute of the computational cost of propagating gradients backward through time in the recurrent layer. The n-back task requires the gradient to flow one more timestep backward through the recurrent dynamics with each additional level of difficulty. The output layer receives a direct gradient signal at every time step, regardless of task difficulty, so the optimizer concentrates its marginal learning effort here where the gradient signal is most reliable. Once the network has learned to encode the input characters into its hidden state, further tweaking of the input weights contributes little to solving a more difficult memory problem.

**Figure 6:**
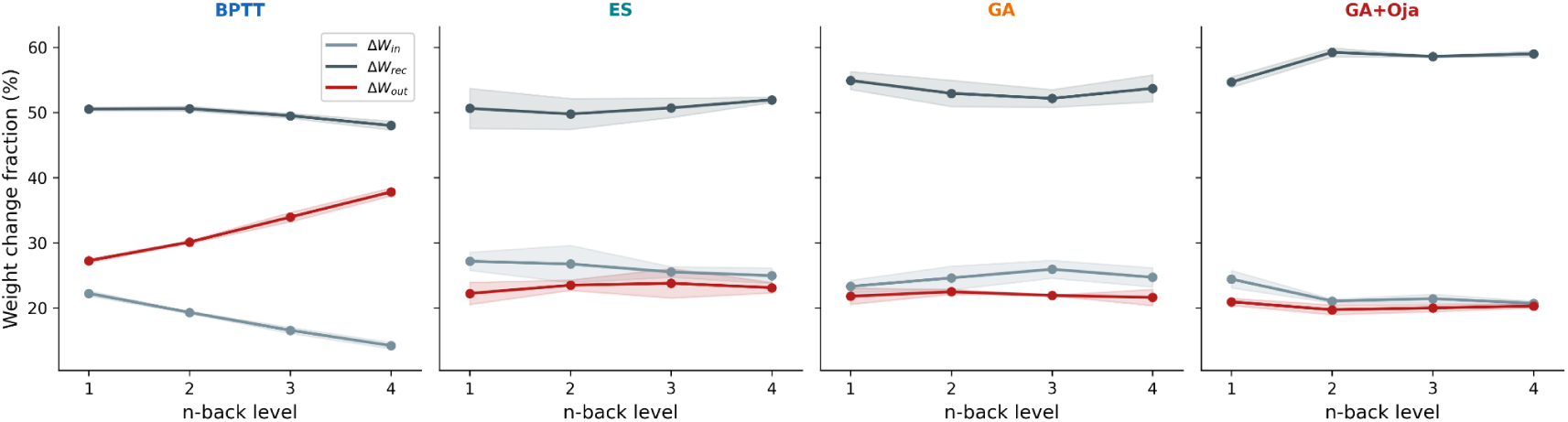
Per-layer weight-change fractions across methods and n-back levels (N = 32 neurons; 10 seeds; mean ± std). BPTT shows a monotonically increasing W_out_ fraction with task difficulty (27.4% → 37.9%; Spearman ρ = +0.969, p_adj_ < 0.001) and a corresponding decrease in W_in_, while W_rec_ remains stable. EA methods show no consistent directional trend across sizes: ES trends slightly negative at N = 64 and N = 128 (ρ = −0.558 and −0.360), GA is weakly positive at N = 32 only (ρ = +0.492), and GA+Oja is largely flat, none approaching BPTT’s effect size.

**Figure 7:**
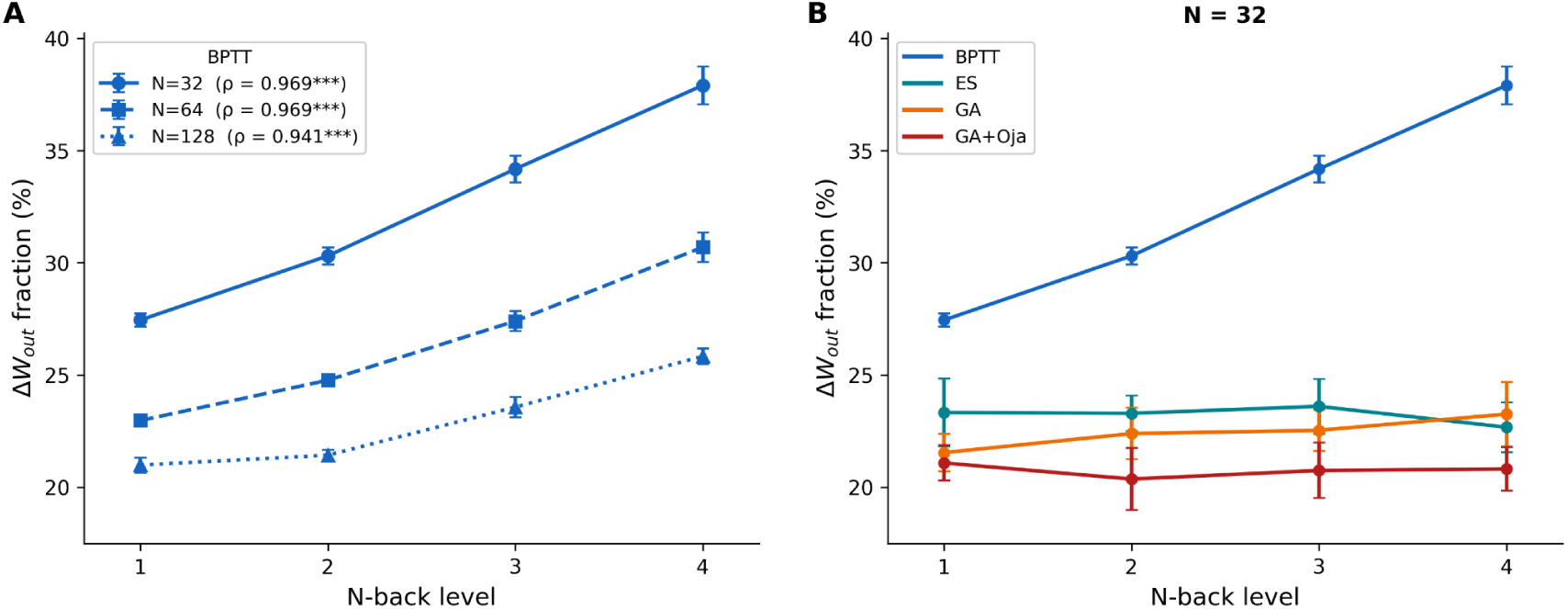
W_out_ weight change fraction vs. n-back level. **A:** BPTT across network sizes (N = 32/64/128), showing a consistent increase with task difficulty at all sizes (Spearman ρ≥0.941, p_adj_<0.001 for all N). **B:** All methods at N = 32, showing BPTT’s rising trend (27.4%→37.9%) against EA methods with inconsistent, small-magnitude trends.

### 3.4 Total Weight Change Magnitude

Total weight change magnitude (Σ ∥Δ*W*∥*_F_* across all layers) differed systematically across methods (Fig 8). BPTT maintained a relatively constant total weight change magnitude (∼12-13) regardless of n-back level, suggesting that it found solutions of similar distance from initialization across difficulty levels. GA and GA+Oja showed increasing total weight change with difficulty, from ∼9 at 1-back to ∼20-22 at 4-back, indicating that the evolutionary search moved further from initialization as the task became harder. ES showed the smallest total changes (∼5-7), consistent with its fine-grained gradient estimation approach that makes small, directed updates.

**Figure 8:**
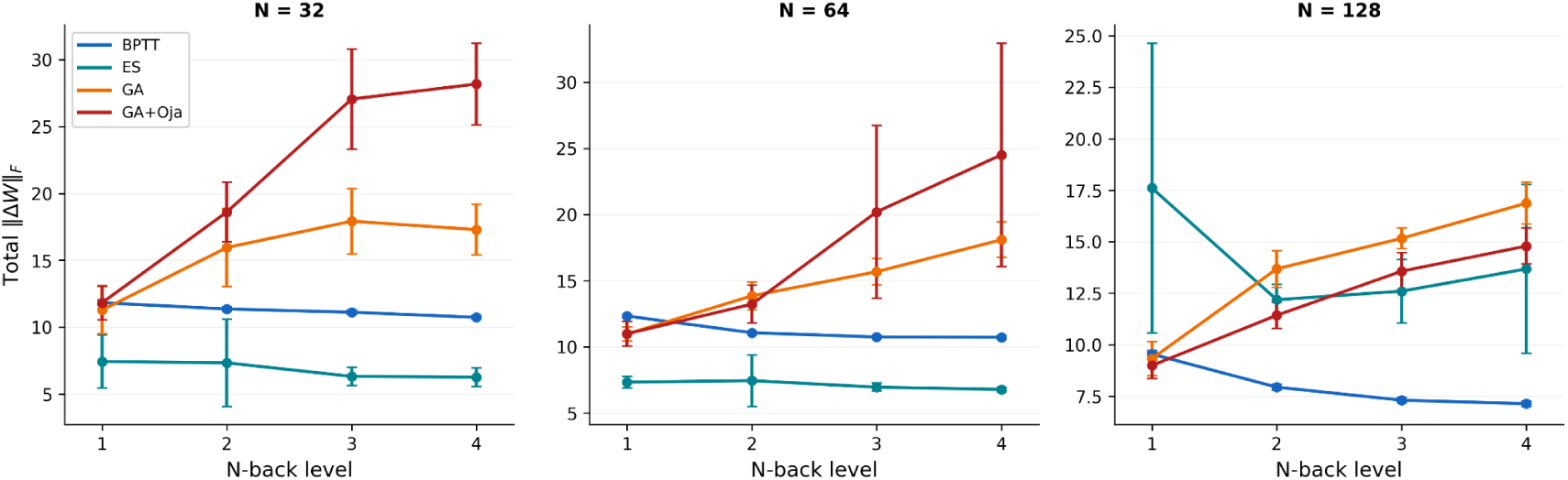
Total weight change magnitude (Σ∥ΔW∥*_F_* across all layers) vs. n-back level (N = 32, 64, 128 neurons; 10 seeds; mean ± std). BPTT maintains roughly constant total change across difficulty levels; GA and GA+Oja increase with difficulty, indicating evolutionary search moves further from initialization at harder tasks; ES consistently shows the smallest total changes, reflecting its directed gradient-estimation approach.

### 3.3 Learning Dynamics

Learning curves revealed qualitative differences in convergence dynamics across methods (Fig 9). BPTT converged rapidly, typically reaching near-perfect accuracy within 50–200 iterations at *N* = 32. ES showed smoother but slower convergence, plateauing over 200–400 generations. GA and GA+Oja showed noisier convergence with greater variance across seeds, reflecting the stochastic nature of population-based search without a gradient estimate. At higher n-back levels, the evolutionary methods’ learning curves often plateaued well below 100% at small network sizes, indicating that the performance gap at those sizes is not a matter of insufficient training time but reflects the difficulty of temporal credit assignment at small scales.

**Figure 9:**
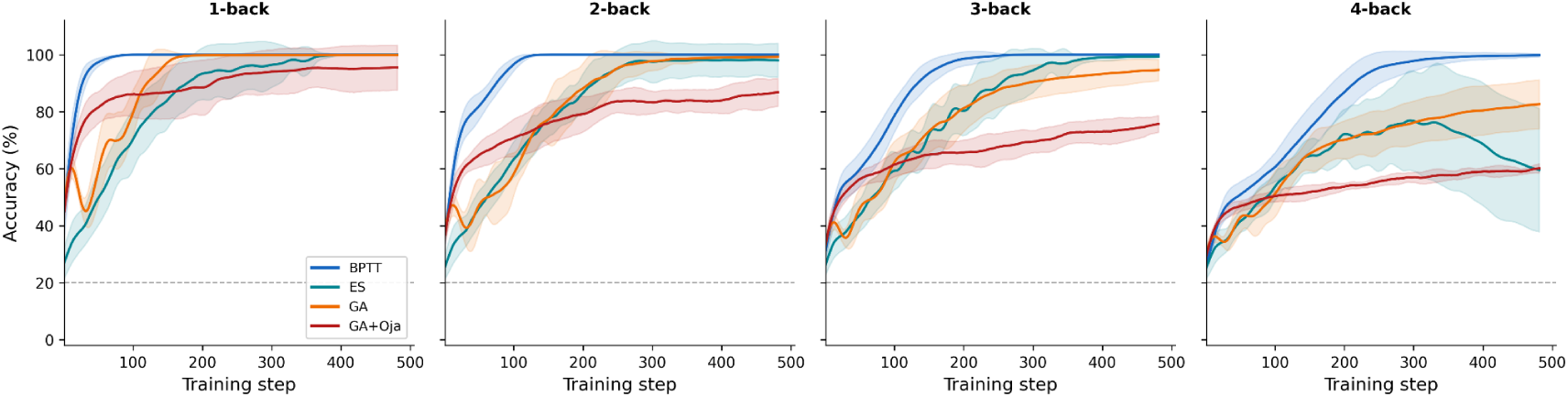
Learning curves for all methods at n-back levels 1-4 (N = 32 neurons; 10 seeds; mean ± std shaded). BPTT converges within 50-200 iterations. ES converges more slowly with higher variance. GA and GA+Oja show noisier trajectories; at n-back 3-4, EA methods plateau below 100%, confirming the performance gap reflects optimization constraints rather than insufficient training time.

### 3.6 Network Activity Dimensionality

The low effective rank of BPTT’s *W*_rec_ predicts the hidden state activity it generates should also occupy a lower-dimensional space. To test this, we ran PCA on the hidden state trajectories of trained networks and measured how many principal components were required to capture 90% of variance across trials.

This prediction was confirmed across all model and task configurations. At *N* = 32, BPTT networks required 5-8 principal components to capture 90% of hidden state variance on the n-back task (increasing slightly with n-back level), whereas ES, GA, and GA+Oja required 11-18 PCs. Cumulative variance curves for BPTT reached 90% rapidly with just a few components, while EA methods showed slower, more distributed variance accumulation consistent with higher-dimensional dynamics (Fig 10). At *N* = 128, BPTT required 11-13 PCs compared to 20-37 for EA methods on the n-back task. On the two-joint arm task at *N* = 32, BPTT required 5 PCS versus 13-16 for EA methods.

**Figure 10:**
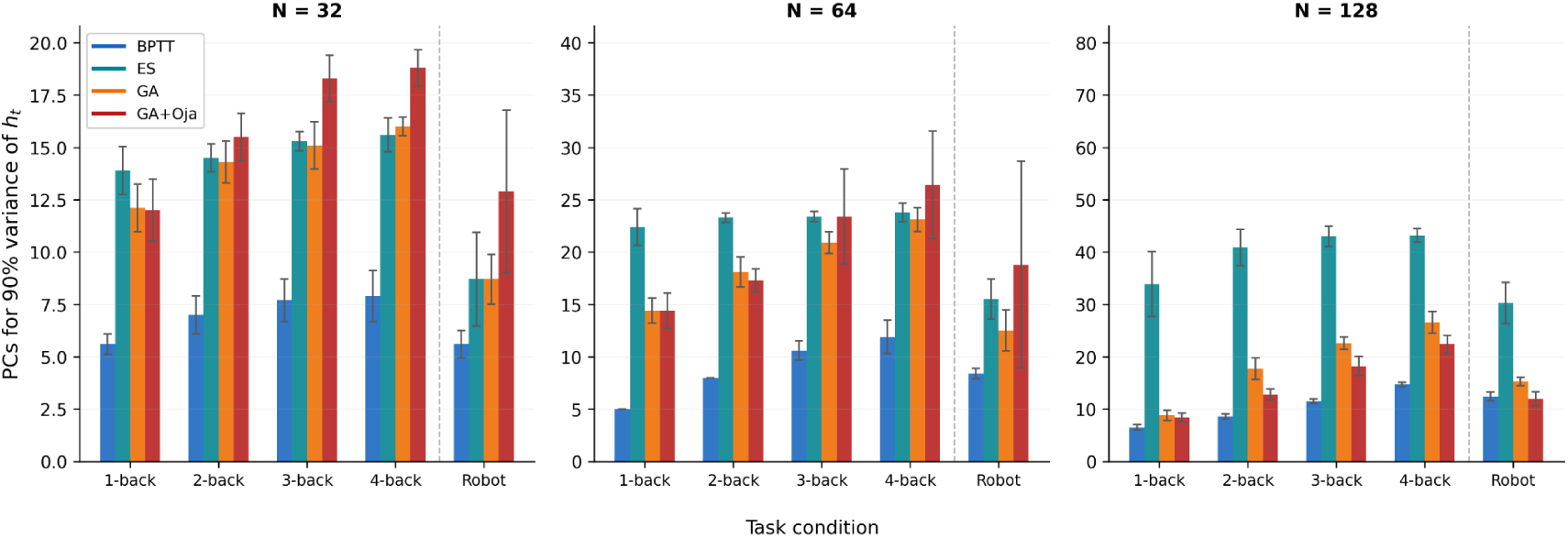
Number of principal components required to explain 90% of hidden-state variance (h_t) by training method, task condition, and network size (mean ± std across 10 networks per condition). BPTT consistently requires fewer components than all three EA methods at every n-back level and network size tested. Activity dimensionality increases slightly with increasing n-back level for all methods.

The low-dimensional structure of BPTT’s activity is directly observable from the geometry of hidden-state population activity in PC space. BPTT-trained networks organize activity into five discrete clusters and capture 57% of variance in 3 dimensions when projecting hidden states from 200 trials onto the top 3 principal components (Fig 11, top). These five clusters correspond to the five letters (A-E) being maintained in working memory. EA methods spread activity across a much larger volume with no tight clustering in the top 3 PCs (ES 26%, GA 27%, GA+Oja 35% variance in top 3 PCs). EA networks may also have specific symbol structure, but these representations are not visible in a 3 PC projection.

**Figure 11:**
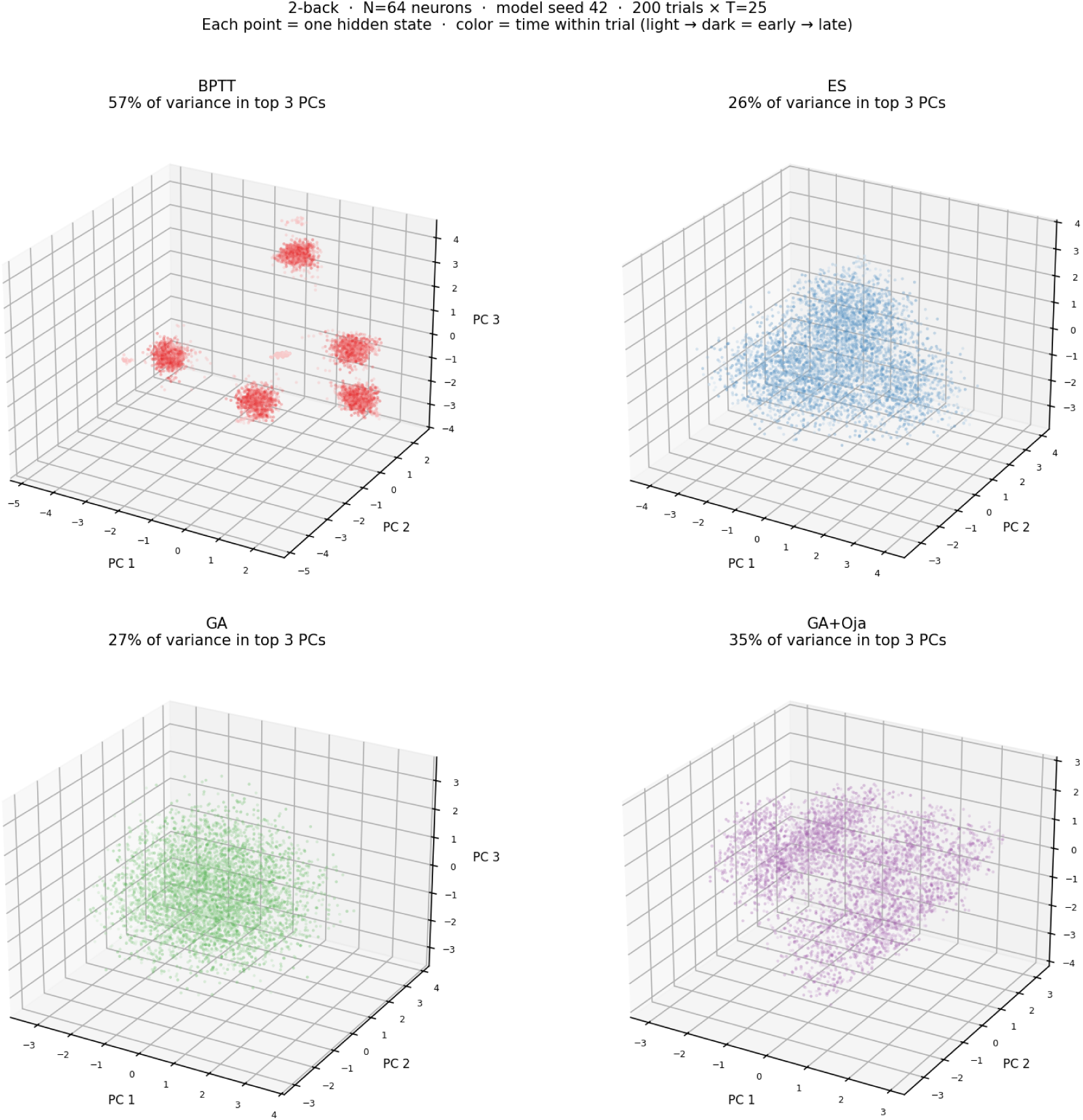
Hidden-state population in PC space during the 2-back task (N = 64, model seed 42). Each panel shows the state-space clouds of all hidden states from 200 trials projected onto the top 3 PCs (each point = one hidden state at one timestep). Trajectory opacity encodes time (lighter = earlier). BPTT organizes activity into five discrete clusters, corresponding to the five letters being maintained in working memory. EA methods distribute activity diffusely across a larger volume of PC space. No clusters are obvious in the top 3 PCs, suggesting symbol-specific structure may exist in higher dimensions.

BPTT’s activity dimensionality was lower than its *W*_rec_ effective rank at matched network sizes. For example, ∼5-8 PCs from activity compared to effective rank of ∼8-10 from weights at *N* = 32. Effective rank measures the dimensionality of the weight matrix itself, so a higher rank is expected when compared to activity PCA, which measures the dimensionality of the subspace actually visited during task performance.

### 3.4 Two-joint Arm Task

The two-joint arm endpoint prediction task differs from the n-back task in a fundamental way: it is not a working memory task. There is no discrete representation to store and retrieve; instead, the network must continuously transform a stream of angular velocity inputs into endpoint position predictions at every timestep. The output at time t depends on the cumulative integral of all preceding inputs (*φ*(t) = Δt · Σ *ω*(t*^′^*)), meaning the recurrent dynamics must implement an ongoing integration and coordinate transformation rather than a store-and-recall operation. This distinction is critical for interpreting the connectivity signatures: any differences from the n-back pattern reflect how the training method adapts to a task that demands continuous dynamical computation rather than memory maintenance.

The connectivity changes of the trained arm task network are different from those of the n-back task network. At *N* = 32, BPTT allocated 63.3% of weight change to *W*_rec_ on the arm task compared to 49.5% pooled across n-back levels (U = 0, *p*_adj_ < 0.001, *r*_rb_ = 1.000). *W*_out_ fell correspondingly from 32.5% (n-back) to 19.2% (arm task). This reversal (*W*_out_-heavy for working memory, *W*_rec_-heavy for continuous dynamics) was consistent across all network sizes (*r*_rb_ = 1.000 at *N* = 32, 64, and 128; all *p*_adj_ < 0.001).

On the arm task, BPTT and the EA methods converged on the same layer-allocation strategy, with both shifting weight change toward *W*_rec_. Crucially, all EA methods also showed significant increases in the fraction of weight changes in *W*_rec_ (frac_rec_) on the arm task (all *p*_adj_ < 0.001, *r*_rb_ = 1.000 at *N* = 32 for GA and GA+Oja; *r*_rb_ = 0.975 for ES). However, the magnitude and interpretation differ. At *N* = 32, the delta in frac_rec_ (arm − n-back) was +0.138 ± 0.017 for BPTT versus +0.144 ± 0.012 for GA (U = 38, *p*_adj_ = 1.000, not significantly different) and +0.085 ± 0.023 for ES (U = 97, *p*_adj_ = 0.009, *r*_rb_ = −0.940). At *N* = 64, BPTT’s shift (+0.142) was significantly larger than GA (+0.108, *p*_adj_ = 0.005) and GA+Oja (+0.101, *p*_adj_ = 0.005). At *N* = 128, BPTT’s shift (+0.092) is comparable to ES (+0.097; not significantly different) and significantly larger than GA+Oja.

BPTT’s recurrent rank increased overall on the arm task relative to n-back, and the gap between recurrent ranks when trained with BPTT vs. EA shrank. For the arm task, effective rank at *N* = 32 was slightly higher for EA, just as it was for the n-back task: BPTT at 15.2 ± 0.4, versus EA methods at 16.4-16.8. At *N* = 128, this gap nearly closed: BPTT 65.2 ± 0.4, EA methods 65.4-65.6. The effective rank for BPTT on the arm task was higher than on the n-back task at *N* = 32 (15.2 vs. 8.4-10.2), suggesting that continuous sensorimotor computation can require a richer recurrent subspace than discrete working-memory maintenance.

All training methods shifted weight changes toward *W*_rec_ on the arm task, but for BPTT the shift was especially large because BPTT-trained models started with heavier *W*_out_ allocation on the n-back task. The EA methods, particularly GA, allocated changes heavily to *W*_rec_ on both tasks: GA at *N* = 32 reached 67.0% frac_rec_ on the arm task and 50.5 – 54.5% across n-back levels, with *W*_out_ staying in a narrow band of 21.5 – 23.3%. In other words, GA’s weight change allocations do respond to task type, but they do not exhibit the *W*_out_ concentration that BPTT shows with increasing n-back task difficulty.

We discuss these task and training method differences further in section 4.4.

### 3.5 GA+Oja: Decomposing Evolutionary and Plasticity Contributions

GA+Oja consistently underperformed plain GA in task accuracy across all network sizes (Mann–Whitney U on pooled n-back 1–4: all *p*_adj_ < 0.001; *r*_rb_ ranged from −0.63 at *N* = 32 to −1.00 at *N* = 128). It produced no significant improvement in *W*_rec_ effective rank at any tested size (all *p*_adj_ > 0.17; a weakly higher median at *N* = 32 did not survive correction for multiple comparisons). Adding within-lifetime Hebbian plasticity to GA therefore reliably reduced task performance without any compensating structural benefit.

**Figure 12:**
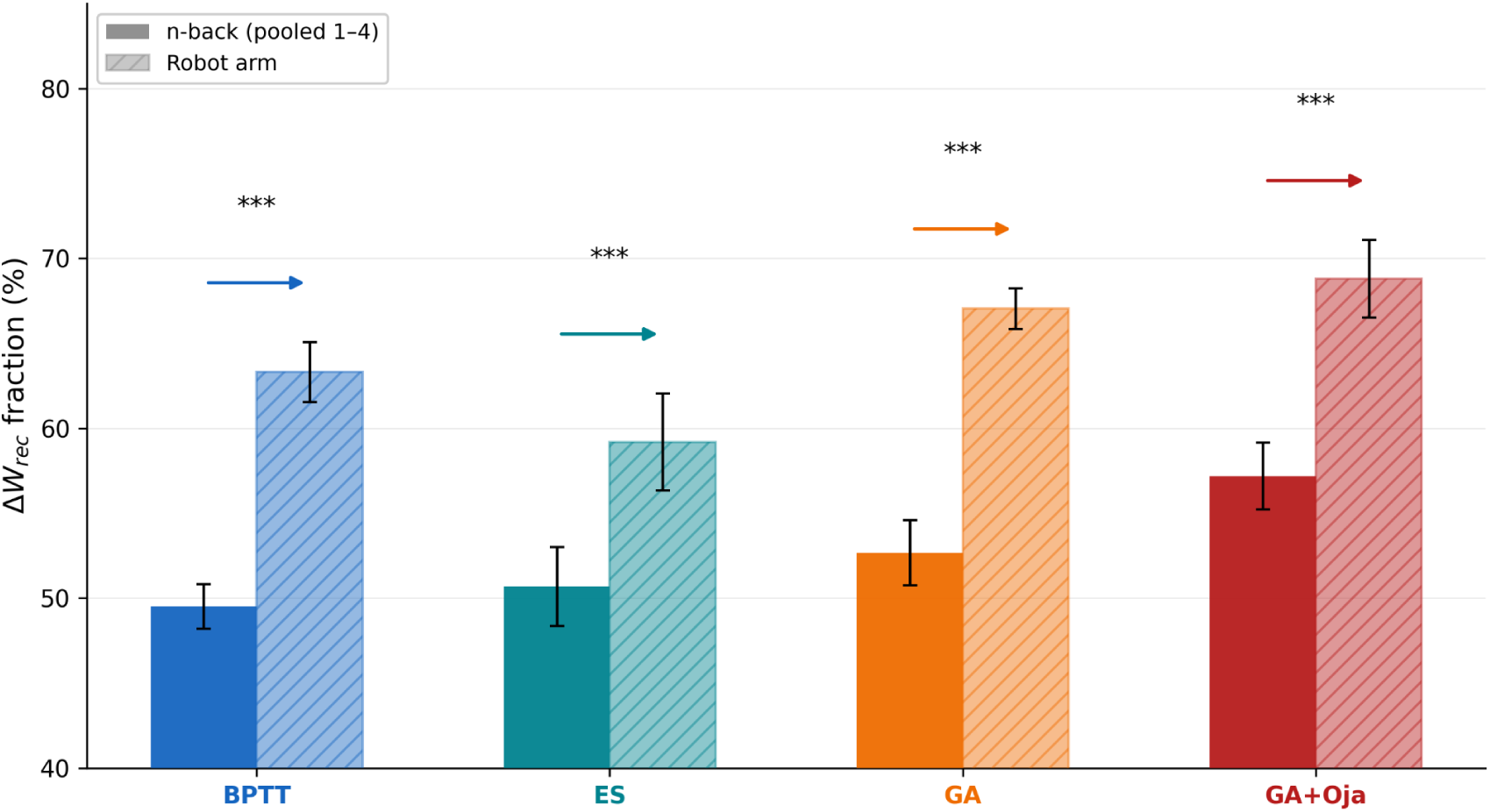
Recurrent weight change fraction (frac_Δ__Wrec_) for n-back (pooled levels 1-4) vs. two-joint arm task, by method (N = 32 neurons; 10 seeds; mean ± std). All methods shift toward greater recurrent weight change on the arm task (all p_adj_<0.001, r_rb_=1.000). BPTT’s distinction is its higher W_out_ emphasis on the n-back task (up to 37.9% at 4-back), making the cross-task contrast largest in absolute terms. EA methods also shift substantially, with GA’s frac_rec_ on the arm task (67.0%) comparable to BPTT’s (63.3%).

Despite this accuracy cost, GA+Oja produced a structurally interpretable decomposition of connectivity change. Oja’s rule modified only *W*_rec_ by construction, and these within-trial changes were fully discarded after each trial with no Lamarckian inheritance, so the genotype’s *W*_rec_ was preserved unchanged for the next generation. The evolutionary contribution 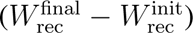 was distributed across all three layers, while the plasticity contribution 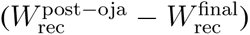 was confined entirely to *W*_rec_.

The magnitude of the Oja contribution was itself under evolutionary control. At *N* = 32 on the two-joint arm task, 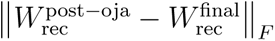 ranged from 0.17 to 15.4 across seeds, depending on the evolved *η* and *w*_max_ parameters. When the GA found plasticity parameters in a productive regime (*w*_max_ ≈ 0.10), the Oja contribution was moderate and the network retained reasonable task performance. Outside of the productive parameter regime, the Oja contribution was sometimes disruptive: large within-trial weight changes degraded the recurrent dynamics the GA had evolved. This sensitivity to plasticity parameter values, combined with the consistent accuracy cost relative to plain GA, shows that Oja’s unsupervised Hebbian rule creates a conflict rather than a contribution to task performance, which we discuss in section 4.5.

## 4 Discussion

Our results demonstrate that training algorithms leave characteristic structural signatures on recurrent network connectivity that are largely independent of task performance. BPTT consistently produces low effective rank in *W*_rec_, reallocates weight change toward the output layer as task difficulty grows, and confines hidden-state activity to a low-dimensional manifold during task performance. Evolutionary methods, in contrast, maintain high effective rank, show no comparable difficulty-dependent reallocation toward *W*_out_, and generate higher-dimensional activity dynamics. These signatures are present even when methods achieve comparable accuracy, indicating that they reflect properties of the optimization algorithm rather than the computational requirements of the task itself.

### 4.1 BPTT’s Low-Rank Bias and Its Dynamical Consequences

BPTT’s low-rank recurrent weights align with a growing body of theoretical work on the implicit biases of gradient descent. In overparameterized networks, gradient-based optimization tends to converge on solutions with low-rank weight matrices even without explicit regularization (Gunasekar et al., 2017; Arora et al., 2019). Our results confirm this low-rank bias and extend it: the low-rank structure of *W*_rec_ is not an isolated property but predicts a corresponding reduction in hidden-state dimensionality during task performance. BPTT produces networks that are low-rank in both structure and dynamics.

The untrained recurrent weights are high rank at initialization (16.6 of a possible 32 at *N* = 32), whereas EAs hold high rank while matching BPTT accuracy at 1- and 2-back. The rank difference is the critical result: the same computation can be carried out with either high- or low-dimensional recurrent connectivity, and which one emerges is determined by the training algorithm. The low effective rank of BPTT-trained networks is thus an algorithmic signature rather than a feature of the computationally optimal solution.

A second line of evidence for BPTT’s low-dimensional activity comes from the geometry of hidden-state trajectories in PC space. BPTT-trained networks (64 neurons, 2-back task) organize activity into five discrete clusters that together capture 57% of total variance in the top 3 PCs. These clusters correspond directly to the five symbols (A–E) maintained in working memory: BPTT has shaped the recurrent state space into five distinct regions, and activity settles into the region appropriate to whichever letter is currently being held. This symbol-by-region structure is the geometric expression of BPTT’s gradient-driven solution, whereas EA methods distribute activity diffusely across a larger volume with no such clustering visible in 3 PCs, yet achieve comparable task performance at matched difficulty. High-dimensional recurrent dynamics are therefore not an obstacle to solving working memory tasks, nor are low-dimensional dynamics an explanation for how such tasks must be solved; they are simply different solution regimes that are uncovered by different training methods.

Low-dimensional recurrent dynamics have been widely observed in the computational neuroscience literature: BPTT-trained RNNs consistently exhibit low-dimensional dynamics on motor tasks and working memory tasks. This has been interpreted as evidence that neural circuits operate in low-dimensional spaces (Mante et al., 2013; Sussillo & Barak, 2013; Cueva et al. 2020, among others). Our results suggest that this interpretation requires caution on two fronts. On the modeling side, we show the same tasks can be solved by networks operating in substantially different dimensional regimes: EAs find higher-dimensional solutions than BPTT and the choice between the two is determined by the training algorithm. Thus, the low-dimensional dynamics in BPTT-trained models reflect the optimizer’s inductive bias.

From the experimental side, low-dimensionality in neural population recordings could be an added artifact of over-training on overly stereotyped tasks. Limited behavioral variables may act as an upper bound on recoverable neural dimensionality, potentially masking high-dimensional population codes (Gao & Ganguli, 2015). Large-scale recordings of spontaneous activity show far higher-dimensional structure than task-focused recordings measure (Stringer et al., 2019), and pre-frontal cortex exhibits high-dimensional activity tied to nonlinear mixed selectivity to enable flexible behaviors (Rigotti et al., 2013).

Of course, the literature itself is full of nuance. Use of discrete object categories lead to low-dimensional memory representations in prefrontal cortex (Sakelliadou et al. 2026), while in the hippocampus each episodic memory is bound with a specific, high-dimensional “barcode” which arises from strong recurrence in RNN models (Fang et al. 2024). Theoretical work suggests that low-dimensional dynamics support noise suppression, leading to a longer working memory capacity (Dinc et al. 2026), while a collapse in prefrontal cortex activity dimensionality is linked to greater error in animal performance (Rigotti et al. 2013). One proposed explanation has been that low-dimensional dynamics occur as the stable results of high-dimensional active computations (Singer & Lazar 2016). Researchers have found that visual cortical activity in response to drifting gratings can be reduced to low-dimensional latents, while the responses to natural images cannot (Schmutz et al. 2026). Motor cortex is where the evidence for low-dimensional dynamics has historically been strongest, but more recent work adds inflections: motor actions with different kinematics may require different levels of dimensionality–i.e. grasping is high-dimensional while reaching is low (Suresh et al. 2020), and even for reaching, multiple different planes are used for distinct reaches (Sabatini & Kaufman 2024).

Taken together, these considerations caution against treating low-dimensionality as a general principle of how the brain solves tasks. The standard modeling pipeline (gradient-trained RNNs) push toward low-dimensional measurements, while the experimental literature suggests a high degree of nuance and specificity. When low-dimensional dynamics from trained RNNs are interpreted as what the brain is optimally doing, these features may in fact have been introduced by the optimizer. Our work shows that high-dimensional solutions are viable and reachable with alternate training methods, and motivates reconsideration of whether the brain may solve cognitive tasks in higher-dimensional regimes than those typically reported from BPTT-trained models.

### 4.2 Global Error Signals

Our comparison of BPTT against gradient-free methods raises the question of whether biological circuits receive information similar to the error signals which form BPTT’s low-rank bias. There is no known biological mechanism for global loss computation or precise temporal credit assignment as there is in BPTT. However, the brain does learn from errors in a temporally structured manner, and a growing body of work investigates whether local neuromodulatory signals, predictive coding frameworks, or local learning rules approximate backpropagation (Lillicrap et al., 2016; Sacramento et al., 2018; Millidge et al., 2022). Though the brain lacks the precise, layer-wise temporally unrolled gradient that BPTT computes, the brain does receive error signals in various forms. Dopaminergic prediction error signals, for example, encode the difference between expected and actual received reward using phasic firing in midbrain dopamine neurons. This process uses a scalar global signal which modulates synaptic plasticity using eligibility traces rather than layer-specific gradient propagation (Frémaux & Gerstner, 2016).

If biological circuits have access to error-like feedback at their readout stage (as suggested by dopaminergic prediction error signals) but lack a mechanism for propagating the feedback precisely through recurrent dynamics across time, then we might expect biological networks to follow an intermediate between our EA and BPTT results. The readout stages may be shaped by precise error feedback, while recurrent dynamics within intermediate regions may remain less precisely compressed, shaped by the need for flexibility and evolutionary and developmental constraints. EA-trained RNNs can then still be used as a plausible model for recurrent neural computations.

### 4.3 BPTT’s Adaptive Layer Allocation and Reservoir Computing

The progressive shift of BPTT’s weight allocation toward *W*_out_ with increasing n-back difficulty reveals that gradient-based training adapts its learning strategy to match the computational demands of the task in a specific and interpretable way. The greater the working-memory demand, the more BPTT leans on the readout. For example, at *N* = 32 and 4-back, BPTT allocates 37.9% of total weight change to *W*_out_, up from 27.4% at 1-back.

The n-back task is fundamentally a working memory task: the network must encode a symbol into a stable internal representation, maintain that representation across a delay, and then decode it at the appropriate time. The gradient must propagate further backward in time to update the weights for tracing the correct symbol with higher n-levels. The recurrent dynamics serve as a maintenance buffer, and the computational challenge at higher n is reading out the correct symbol from an increasingly crowded memory trace. In contrast, the output layer receives a direct gradient signal at every timestep regardless of task difficulty – BPTT responds by concentrating additional learning effort in the output layer when difficulty increases. It is crucial to note that EA and BPTT reach equivalent accuracy at *N* = 64 and *N* = 128 even for high n levels (Table 1). Thus the same working memory task can be effectively solved by two differing strategies: 1) refinement of the readout of recurrent dynamics (under BPTT) or 2) refinement of the recurrent dynamics themselves (under EAs).

This result complements the higher dimensional dynamics of EA-trained networks compared to BPTT-trained networks. High dimensional dynamics carry the benefit of enabling even simple readouts to decode a large set of input-output relations (Rigotti et al. 2013), thus the output layer of EA-trained models does not need to undergo extensive changes to solve the n-back task. Indeed, EAs emphasize a solution that fully utilizes the recurrent layer, whereas BPTT emphasizes the readout.

BPTT’s pattern of concentrating learning in the output layer follows the logic of echo-state networks and reservoir computing (Jaeger, 2001; Maass et al., 2002). A reservoir computer uses a large group of recurrent, randomly connected units (the reservoir) to process incoming temporal data. As inputs arrive, the reservoir’s fixed internal connections generate high-dimensional patterns to encode both the present and recent history of the input. Crucially, only the final readout layer is trained, usually as a simple linear mapping, while recurrent connections within the reservoir remain randomly initialized and unchanged throughout learning (Schuman et al., 2022).

Now, BPTT does not freeze *W*_rec_ in our models entirely. Our results show a trend towards reservoir behavior, in which the difficulty-sensitive work is increasingly absorbed by the output decoder. Likewise, training paradigms that are elaborations upon the strict reservoir framework, such as FORCE learning (Sussillo & Abbott 2009), which can modify recurrent dynamics through feedback, are extremely successful at instilling desired behaviors in RNNs. A practical implication is that in analyses of BPTT-trained models of working memory, decoding should not be regarded as a trivial step of computation. The readout layer can take on a substantial share of the task-sensitive work and should be analyzed alongside the recurrent dynamics.

### 4.2 Cross-Task Comparison: Layer Change Reallocation in the Two-Joint Arm Task

The two-joint arm task presents a fundamentally different computational demand than the n-back task. Instead of storing and retrieving discrete signals, the network must sustain continuous, real-time computation: integrating angular velocities, tracking joint angles, and evaluating the nonlinear forward kinematics at every timestep to estimate endpoint position. In this task, computation unfolds dynamically across the entire trial as opposed to the encode-maintain-decode structure of working memory tasks.

BPTT solves this task by further concentrating learning in *W*_rec_ compared to in the n-back task, consistent across all network sizes. This cross-task difference can either be understood as 1) the gradient concentrates learning where it genuinely improves performance most, or 2) the gradient struggles to fully propagate backwards through time in the recurrent layer for the n-back task. We believe both explanations play a role.

In the n-back task, the main challenge is choosing the correct symbol from memory. Each symbol in a sequence is discrete and independently chosen, so while recurrent activity belies past activity (and thus past symbols), recurrent states are not causally linked through the task itself. In the arm task, however, they are: inputs and outputs are continuous and interrelated in time. One can take this to mean that emphasizing robust readout is more important for the n-back task, while emphasizing appropriate ongoing recurrent dynamics is more important for the arm task. This is consistent with explanation 1: the nature of the arm task leads to more recurrent changes. However, the differing natures of the two tasks also means that it is genuinely harder to make appropriate recurrent changes for the n-back task, consistent with explanation 2: improper gradient propagation leads to more output changes for the n-back task.

In support of explanation 1, EAs as well at BPTT show a preference for *W*_rec_ changes on the arm task: all three EA methods shift with *p*_adj_ ≤ 0.0001 at every network size, and GA’s shift at *N* = 32 is comparable in magnitude to BPTT’s (GA +0.144 vs BPTT +0.138 in frac_rec_; not significantly different). Since both training methods converge on the same layer-wise change allocation solution, this suggests a genuine, performance-based reason to have more *W*_rec_ changes in the arm task relative to the n-back task.

However, what EA methods do not replicate is BPTT’s within-task reallocation with difficulty. On the n-back task, BPTT’s *W*_out_ fraction climbs monotonically from 27.4% at 1-back to 37.9% at 4-back, a 10.5-percentage-point shift. In other words, as gradients need to propagate back further in time, *W*_out_ changes increase. The analogous EA trends are small and inconsistent: at *N* = 32, GA shows a weak positive trend but the shift is only 1.8 percentage points; ES and GA+Oja show no significant trend at *N* = 32; and at larger sizes the direction varies by method (ES trends slightly negative, GA and GA+Oja trend weakly positive or flat). No EA method reproduces BPTT’s difficulty-scaled reallocation toward the readout. These results suggest that a component of BPTT’s cross-task reallocation (*W*_out_-dominant for working memory, *W*_rec_-dominant for continuous dynamics) may arise from the nature of gradient-based training and not only from the nature of the computations.

### 4.3 EA Performance and Scaling

GA+Oja was designed following the biological intuition of two processes that shape neural circuit connectivity on different timescales. Evolution shapes the initial wiring over generations, whereas within-lifetime learning refines connections from experience. GA+Oja implements the separation explicitly: GA evolved the initial weights across generations and Oja’s unsupervised Hebbian rule modified *W*_rec_ within each trial. Evolutionary weight changes were distributed across each layer, and Hebbian plasticity changes were confined to *W*_rec_. GA+Oja consistently underperformed plain GA across all network sizes and difficulty levels.

When the GA found productive *η* and *w*_max_ values, the Oja contribution was moderate and the network retained reasonable task performance; when it did not, the Oja contribution was either negligible or disruptive enough to degrade recurrent dynamics the GA had evolved. While on the surface not the result we had anticipated, we believe the negative contribution of Oja’s rule in fact supports our finding of higher-dimensional solutions under EAs.

Oja’s rule, like many Hebbian plasticity rules by design, promotes greater correlation in population activity within bounds. Neurons that are co-active will have their synapses further strengthened, so that their activities become even more correlated over time, thus reducing the overall dimensionality of population dynamics. By contrast, we have found that EAs tend to arrive at high-performing, high-dimensional dynamic solutions. The within-lifetime insertion of Oja’s rule disrupts this dimensionality, leading to degradation of performance. Where then does this leave us, considering the reality that neural connections are changed in productive ways by both evolution and within-lifetime local plasticity? Oja’s rule is a simplification of local plasticity rules observed in the brain, where anti-Hebbian forces work alongside complex Hebbian ones. Future works on hybrid evolutionary-plasticity approaches should account for this as best they can.

### 4.4 Implications for Computational Neuroscience

The central implication of our findings for computational neuroscience is that structural analyses of trained RNNs must account for the training algorithm as a potential confound. When researchers train an RNN on a cognitive task and then analyze its connectivity, dynamics, or representations, they are studying a composite of task-determined structure and algorithm-determined structure. Our results show that these two contributions can be disentangled by comparing networks trained with different algorithms on the same task.

Concretely, our findings suggest that low-dimensional recurrent dynamics observed in BPTT-trained RNNs may reflect the optimization algorithm’s implicit bias rather than a computational necessity. Studies have shown that low-dimensional dynamics occur in real cortical recordings under many circumstances–this is well-established. However, some studies also follow up with demonstrations that RNNs trained with BPTT exhibit similar low-dimensional dynamics. These models and modeling results should be used with caution, and not be interpreted outright as demonstration of the necessity of low-dimensional computations generally. Studies have also revealed high-dimensional dynamics in cortex under other circumstances, and it is possible that gradient-trained RNNs may underestimate the dimensionality of biological neural computation. Future work in computational neuroscience could directly compare gradient-trained and evolution-trained networks as complementary models of the same underlying system, using their divergences to disentangle which structural and dynamical properties are task-driven versus algorithm-induced.

### 4.5 Limitations and Future Directions

Several limitations qualify our conclusions. First, our networks are small (*N* = 32 to 128) compared to both neurobiological circuits and modern deep learning models. We deliberately used the smallest networks that solve each task to isolate task-relevant principles from capacity-driven bloat. Whether the structural signatures we observe persist at larger scales is an open question. The activity-dimensionality gap between BPTT and EAs and the *W*_out_ reallocation pattern were robust across the sizes we tested, but their scaling behavior in networks with thousands or millions of neurons remains to be determined. Second, we used neurons with continuous activations, not biologically realistic spiking neurons. The non-differentiability of spikes creates additional challenges for gradient-based methods (partially addressed by surrogate gradients) that do not affect evolutionary methods, potentially exacerbating training method biases in spiking models. Third, we did not enforce Dale’s Law, nor specify excitatory and inhibitory populations. Enforcing such constraints would add biological realism but also constrain the solution space in ways that might interact differently with gradient-based versus evolutionary optimization, complicating the interpretation. Fourth, our analysis is limited to two instances of two task classes. Studying connectivity and dynamic signatures across a broader range of cognitive tasks, particularly tasks with contextual shifts and naturalistic stimuli, remains open. We hope that our results in small, simple models and two tasks can 1) act as a springboard for future explorations with more realism and diversity while 2) remaining as a proof-of-concept that similar exterior results can be achieved with differing interior processes (Beer et al. 2024).

Finally, at *N* = 32 on harder n-back levels, GA and ES did not reach BPTT’s near-perfect accuracy even under the dimension-aware scaling described in the Methods (GA 82.8% at 4-back, ES 58.6% at 4-back). Connectivity comparisons at these conditions are therefore between networks at different performance levels. We emphasize, however, that the effective-rank and activity-dimensionality differences are present at 1-back and 2-back where all methods except GA+Oja achieve comparable accuracy, and the differences persist at *N* = 64 and *N* = 128 where EA methods also reach near-perfect accuracy. The algorithmic signatures are therefore not an artifact of performance mismatch, since they hold when performance is matched.

Based on the present limitations, several extensions of this work would strengthen and broaden our conclusions. First, the evolving-connectivity framework of Wang et al. 2023, in which connection probabilities rather than weights are evolved, could be compared against our weight-evolution approach to determine whether probabilistic connectivity evolution produces yet another distinct class of structural signature. Second, extending the analysis to Dale’s-Law-constrained and spiking networks would test whether the BPTT–EA dimensionality gap persists under more biologically realistic architectures, or whether the gap narrows once the solution space itself is biologically constrained. Third, applying our framework to a broader task range, including context-dependent decision-making, continuous-time motor planning, and multi-step reasoning, would reveal whether BPTT’s adaptive *W*_out_ concentration and the EA methods’ high-rank tendencies are universal signatures or task-family-specific. Fifth, extending our study to gradient-based methods outside of standard BPTT would lend clarity to the applicable extent of our results–to gradient-based methods generally, or only to BPTT. Finally, the most tedious and powerful extensions are to revisit the literature of gradient-trained RNNs and test the same models and tasks with gradient-free training methods, and to apply this dual approach when creating models of new cortical experiments in the future. We suspect the description of real cortical computation is simultaneously more complex and more elegant than just “high-dimensional” or “low-dimensional” alone can capture, and we encourage modelers to question the limitations that singular training methods can place on our understanding.

